# Asynchronous subunit transitions precede acetylcholine receptor activation

**DOI:** 10.1101/2024.12.02.626350

**Authors:** Mackenzie J. Thompson, Christian J.G. Tessier, Anna Ananchenko, Johnathon R. Emlaw, François Dehez, Eleftherios Zarkadas, Corrie J.B. daCosta, Hugues Nury, John E. Baenziger

## Abstract

Rapid communication at synapses is facilitated by postsynaptic receptors, which convert a chemical signal into an electrical response. In the case of ligand-gated ion channels, agonist binding triggers rapid transition through a series of intermediate states leading to a transient open-pore conformation. These transitions are usually framed in terms of a mechanism where agonist binding and channel activation are separate events. Here, we collect cryo-EM images over a range of agonist concentrations to define structures of the muscle-type nicotinic acetylcholine receptor in unliganded, mono-liganded, and di-liganded states. We show that agonist binding to a single agonist site stabilizes an intermediate state where an entire principal agonist-binding subunit has transitioned to an active-like conformation, while the other unoccupied principal subunit remains inactive, albeit poised for activation. Binding of agonist to the second agonist site fully activates the remaining subunits leading to hydration of the ion pore. Uniting this cryo-EM derived intermediate structure with single-channel recordings leads to a model where individual acetylcholine receptor subunits asynchronously undergo conformational transitions, and thus a sequential activation mechanism that has implications for the entire superfamily of pentameric ligand-gated ion channels.

## Main

Communication at the neuromuscular junction relies on the activation of a pentameric ligand-gated ion channel (pLGIC), the nicotinic acetylcholine receptor (nAChR). More than sixty years ago, del Castillo and Katz modelled nAChR activation as a multistep process where the neurotransmitter, acetylcholine (ACh), reversibly binds to an inactive/closed receptor, which when bound undergoes a conformational change to an active/open conformation^1^. As it became clear that the nAChR has two ACh binding sites, the scheme was expanded to include two separate binding events, as well as openings from both mono- and di-liganded receptors^2^. Continued improvements in the recording and analysis of single-channel electrophysiological data has led to increasingly complex models of nAChR activation, with current schemes including one or more pre-open intermediate states, deemed ‘flipped’ or ‘primed’^3–5^. Although the whole cell response of the muscle-type nAChR to ACh is governed primarily by di-liganded activation, both disease-causing mutations and various ligands alter pLGIC function by modulating the rates of the transitions between these intermediate pre-open states, underscoring their importance to synaptic communication^3–6^.

Despite their importance, intermediate pre-open states have remained largely phenomenological, only ever being inferred from single-channel recordings^3,4,7^. The majority of available nAChR (and related pLGIC) structures represent the most stable, and thus most populated, ‘end’ states. Kinetic analyses predict that these intermediate states are short-lived, so their fleeting lifetimes are difficult to detect by equilibrium methods, especially at the extremes of ligand concentration used in most structural studies. As a result, except for a few notable exceptions^4,8–14^, little insight into the structures of intermediate pre-open states has been gleaned to date.

To gain structural insight into the nAChR activation mechanism, and specifically intermediate pre-open states, we first established an ACh concentration-dependent ‘conformational response’ for the muscle-type *Torpedo* nAChR reconstituted into lipid nanodiscs using the probe, ethidium bromide (Fig. 1a, discussed below). We then recorded cryo-EM datasets at ACh concentrations spanning this conformational response, with the resulting reconstructions (Fig. 1b-d) showing that ACh binding to a single agonist site, in this case the site located at the interface between the principal α_γ_ and complementary γ-subunit, favours a conformation where the entire α_γ_-subunit has transitioned to an active conformation, which in turn renders the remainder of the transmembrane domain poised for full activation. Meanwhile, the second unoccupied agonist site, located at the interface between the principal α_δ_ and complementary δ-subunit, remains in a resting state-like conformation (Fig. 1c). Finally, we use these structures to inform a simple and plausible kinetic scheme that both reconciles the observed single-channel activity of the human adult muscle-type nAChR and predicts the prevalence of the intermediate pre-open state revealed by cryo-EM. Together, our data constitute direct structural and functional evidence for asynchronous subunit transitions preceding muscle-type nAChR activation.

**Figure 1.**
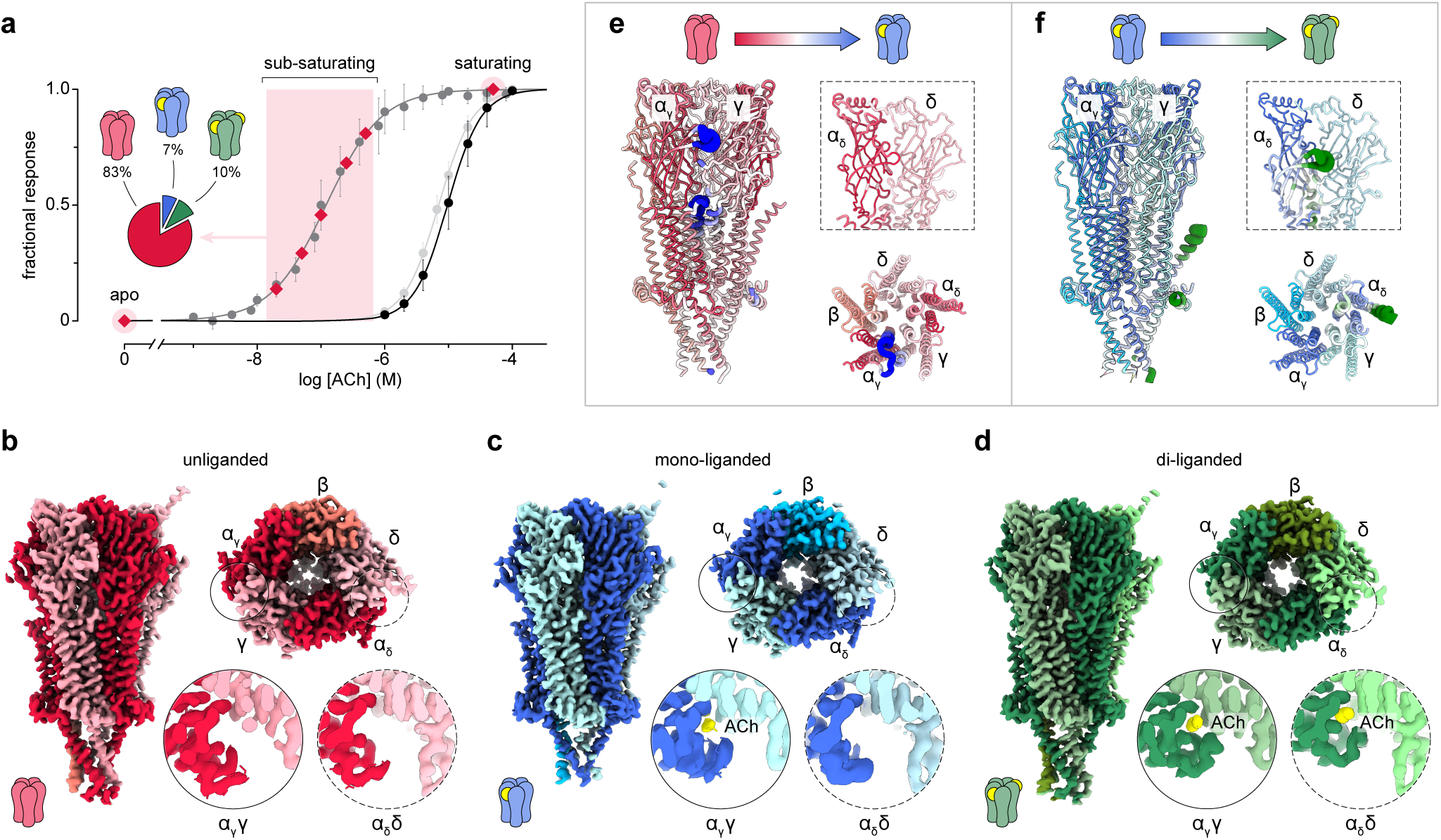
Unliganded, mono-liganded, and di-liganded states. **(a)** Fractional response of the *Torpedo* and human adult muscle-type nAChR to acetylcholine (ACh). The left curve illustrates the ACh-induced binding of ethidium bromide to the MSP2N2-nanodisc reconstituted *Torpedo* nAChR (dark gray). Red diamonds denote the concentrations at which cryo-EM data sets were collected. The pie chart shows the distribution of particles detected in the resulting cryo-EM images recorded at sub-saturating concentrations of ACh (pink box). The two curves on the right show the whole cell current responses measured in *Xenopus* oocytes expressing the *Torpedo* (black) and human adult muscle (light gray) nAChRs. **(b-d)** side and top-down views of cryo-EM reconstructions for **(b)** unliganded (red), **(c)** mono-liganded (blue), and **(d)** di-liganded (right, green) *Torpedo* nAChR, with zoomed in top-down views highlighting loop C capping at both the α_γ_γ (solid) and α_δ_δ (dashed) agonist sites. ACh density shown in yellow. **(e,f)** Cartoon representations highlighting the conformational transition from **(e)** unliganded to mono-liganded and **(f)** mono-liganded to di-liganded states. In each case, a side-view of the nAChR looking at the α_γ_- and γ-subunits color-coded and radius-coded by RMSD to highlight the backbone conformational transition is shown on the left. In both panels, the dashed box on the right shows a side view of the α_δ_δ agonist site, with a top-down view of the transmembrane domain shown below.

### The ACh conformational response

To determine ACh concentrations where pre-open intermediates may be prevalent, we measured the ACh-induced binding of ethidium bromide (Fig. 1a), a conformation-sensitive fluorescent probe that binds with a roughly one-thousand-fold higher affinity to the pore of the desensitized versus the resting state nAChR^15,16^. Ethidium also binds with relatively high, although not quantified, affinity to the open state^17^. We noticed that ACh-induced ethidium binding occurs at ACh concentrations (EC_50_ ∼100 nM) that are roughly one-hundred times lower than those required to induce a whole-cell current response with either the *Torpedo* or human adult muscle-type nAChR (both EC_50_s ∼10 μM)^18,19^. The different EC_50_ values could result from the different membrane environments in which the nAChR is embedded (azolectin-MSP2N2 nanodisc versus *Xenopus* oocyte membranes, respectively). Another possibility is that ACh binding initiates a conformational change in the pore leading to ethidium binding at ACh concentrations well below those that are required to stabilize a fully open state. Indeed, single-channel recordings show that the ACh concentrations used to induce the ethidium conformational response lead to infrequent, isolated channel openings (both mono-liganded and di-liganded), while the higher concentrations required to induce a whole-cell current response stabilize bursts of primarily di-liganded single-channel openings (see below). Hypothesizing that the conformational change detected by ethidium binding relates to an intermediate pre-open state, we set out to solve nAChR structures at ACh concentrations spanning the ethidium response (20, 50, 100, 250, and 500 nM ACh). For comparison, we also recorded data sets both in the absence of ACh and in the presence of saturating 50 μM ACh concentrations.

### Cryo-EM maps of un-, mono-, and di-liganded states

Cryo-EM datasets recorded in the absence of ACh or at saturating ACh concentrations yielded 3D reconstructions corresponding to unliganded and di-liganded states, respectively (Figs. 1b,d and Extended Data Fig. 1). In contrast, datasets collected at sub-saturating ACh concentrations each yielded reconstructions suggestive of a mixture of states in that the distinguishing features between unliganded and di-liganded conformations were not well defined (Extended Data Figs. 2–5). In particular, ACh density in both agonist sites was noticeably weaker than in the di-liganded reconstruction. ACh density was also consistently stronger in the α_γ_γ site than in the α_δ_δ site (Extended Data Fig. 5). We employed a combination of 3D classification and 3D variability analysis^20,21^, testing diverse focused masks for particle sorting around the zones where changes in conformation were most marked (Extended Data Fig. 4). A mixture of unliganded, mono-liganded, and di-liganded conformations were detected in each of the reconstructions, with each mono-liganded conformation containing ACh density exclusively in the α_γ_γ agonist site (Extended Data Fig. 5). We verified, as a control, that it was not possible to detect additional conformations in the reconstructions recorded in either the absence of ACh, or at saturating ACh concentrations. As the quality of every mono-liganded reconstruction obtained from each individual dataset was, by itself, borderline for model building, we combined the datasets collected from all sub-saturating ACh concentrations (20, 50, 100, 250, and 500 nM). After iterative classification, we obtained a final set of particles that yielded a high-quality mono-liganded reconstruction where a β hairpin structure in the α_γ_-subunit, known as ‘loop C’, is capped around ACh in the α_γ_γ agonist site while, at the same time, the corresponding α_δ_ loop C is uncapped with the α_δ_δ site essentially devoid of agonist (Figs. 1c and Extended Data Figs. 5 and 6). Notably, a mono-liganded reconstruction with ACh bound solely to the α_δ_δ agonist site was not detected at any stage of the data processing.

The three reconstructions led to structures corresponding to unliganded (3.2 Å resolution), mono-liganded (3.1 Å resolution), and di-liganded (2.8 Å resolution) states (Extended Data, Table 1). Each exhibits the typical pLGIC architecture, with five subunits arranged pseudo-symmetrically in an α_γ_-β-δ-α_δ_-γ clockwise fashion around a central ion-conducting pore. In each structure, the five subunits each consist of an extracellular domain with ten β-strands, a transmembrane domain with four membrane-spanning α-helices and a short amphipathic α-helix, and an intracellular domain with both an extended α-helix and a substantial disordered stretch (see Fig. 2a).

**Figure 2.**
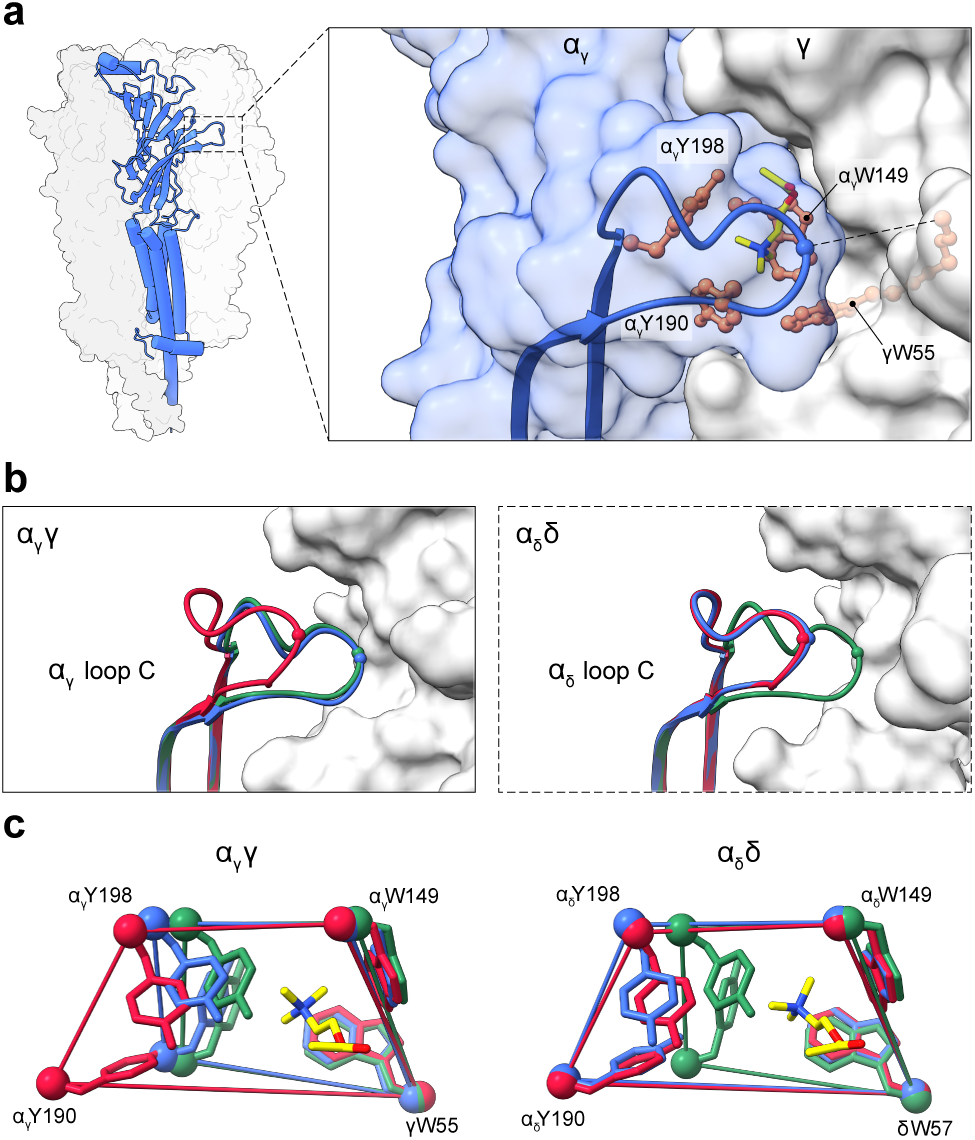
Conformational transitions in the agonist site. **(a)** Side view of the mono-liganded structure with the β, δ, α_δ_ and γ-subunits shown as light surfaces and the α_γ_-subunit as a blue cartoon. The dashed box highlights the region of the α_γ_γ agonist binding site that is zoomed in on the right, with the view rotated to highlight loop C capping. On the right, loop C is shown as a blue cartoon, the side chains that form the α_γ_ aromatic box are shown as orange ball and sticks, and the bound ACh is shown as sticks with color-coded atoms (carbon, yellow; nitrogen, blue; oxygen, red). The dashed line highlights the distance between the C_α_ carbons of α_γ_Cys192 and γGlu57 (or δ Glu59 at the α_δ_δ agonist site), which is used to quantify the degree of loop C capping in the three structures (see text). **(b)** Superimposition of the loop C conformations for the unliganded (red), mono-liganded (blue), and di-liganded structures (green) at both the α_γ_γ (solid box) and α_δ_δ (dashed box) agonist sites. **(c)** Superimposition of the C_α_ carbons (balls) of aromatic box residues (sticks) in the α_γ_γ (left) and α_δ_δ (right) agonist binding sites of the unliganded, mono-liganded, and di-liganded structures.

### The unliganded and di-liganded nAChR

The unliganded nAChR structure with unoccupied solvent accessible agonist sites and a closed, de-wetted channel pore (Figs. 1-4) mirrors those determined previously in the absence of ligand (PDB: 7QKO or 7SMM, RMSDs=0.40 Å and 0.54 Å, respectively), and is similarly assigned to the resting state^22,23^. The di-liganded structure, featuring two bound ACh molecules and an open Leu9’gate, closely resembles those obtained in the presence of carbamylcholine (7QL6 or 7MSR, RMSDs of 0.44 Å and 0.55 Å, respectively) and nicotine (7QL5, RMSD=0.44 Å)^22,23^. Molecular dynamics simulations show the pore of this structure is hydrated and conducts ions, although it quickly collapses to the same de-wetted state observed in previous MD simulations of the carbamylcholine-bound structure (Fig. 4)^23^. Regardless of the assignment to a physiological state, the unliganded and di-liganded structures with closed and open channel pores, respectively, define the global conformational change that leads to both an increase in ACh binding affinity and the opening of the Leu9’ pore gate (see Fig. 6F of ref^23^). This conformational change includes 1) capping of loop C at both α_γ_γ and α_δ_δ agonist sites so that aromatic box residues, αTrp149, αTyr190, αTyr198, and γTrp55/δTrp57, wrap tightly around the quaternary amine of ACh (Fig. 2), 2) upward movement of ‘loop F’ from both complementary γ/δ-subunits to interact with loop C from the adjacent principal α subunit (Extended Data Fig. 7), 3) a global pivoting of the two α subunits toward their adjacent β/γ subunits to facilitate the outward translation of αPro265, within the M2-M3 loop of the transmembrane domain, past αVal46, in the β1-β2 loop of the extracellular domain, a movement that correlates with channel activation (Fig. 3)^19,24^, and 4) an outward tilting and twisting of the pore-lining M2 α-helices away from the central pore axis leading to hydration and thus cation flux through the channel pore (Fig. 4).

**Figure 3.**
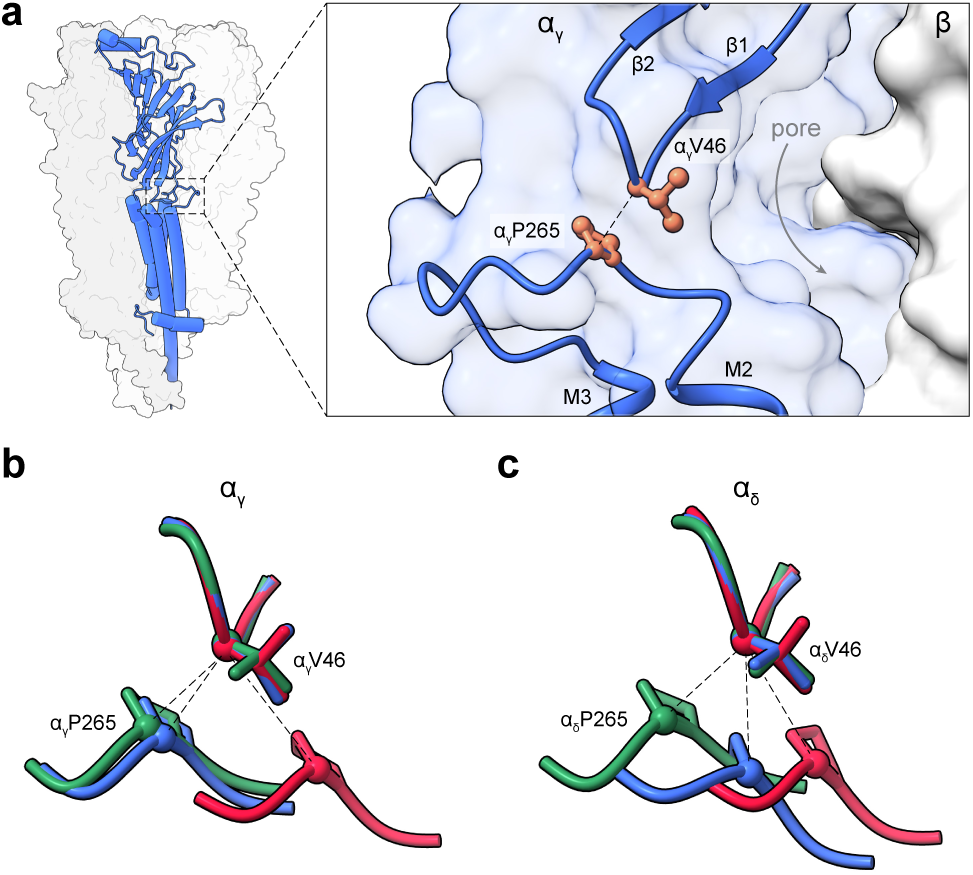
Conformational transitions at the interface between the extracellular and transmembrane domains. **(a)** Side view of the mono-liganded structure with the dashed box highlighting the interface between the extracellular and transmembrane domain of the α_γ_-subunit that is zoomed in on the right. The zoomed in view is rotated to highlight both movements of M2 away from the pore and the relative positions of the β1-β2 and M2-M3 loops from the extracellular and transmembrane domains, respectively, in the unliganded (red), mono-liganded (blue), and di-liganded (green) structures. On the right, the β1-β2 and M2-M3 loops are shown as blue cartoons and the key interface residues, αVal46 and αPro265, are shown as orange ball and stick. **(b)** and **(c)** show superimposed views of the β1-β2 (αVal46) and M2-M3 loops (αPro265) in the unliganded (red), mono-liganded (blue), and di-liganded (green) structures for the α_γ_ and α_δ_-subunits, respectively.

**Figure 4.**
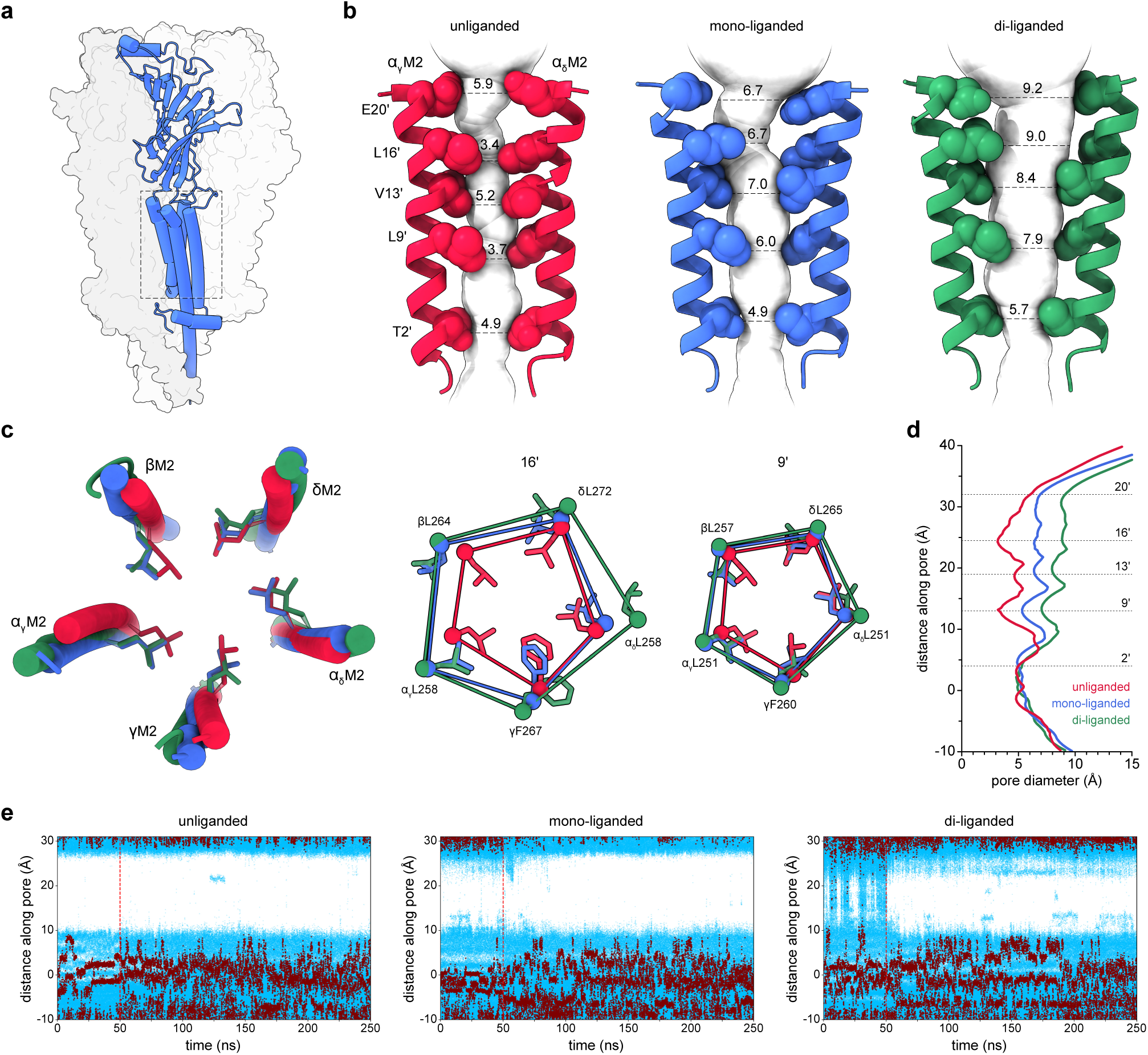
Conformational transitions of the pore. **(a)** Side view of the mono-liganded structure with the dashed box highlighting the transmembrane pore region, with various zoomed in views shown in the remainder of the figure. **(b)** Side views of the α_γ_M2 and α_δ_M2 α-helices. **(c)** Top-down views of both the pore-lining M2 α-helices (cylinders) and the Leu9’ side chains (sticks) from each of the five subunits in the unliganded (red), mono-liganded (blue), and di-liganded (green) states. The C_α_ carbon atoms of the five pore-lining α-helices (spheres connected by imaginary lines) are illustrated at the 16’ and 9’ positions for the three conformations to the right. **(d)** Pore diameters in the three nAChR structures. **(e)** Trajectories showing water (blue) and sodium (red) ion z coordinates of individual particle positions along the channel axis during molecular dynamic simulation runs for the unliganded (left), mono-liganded (middle), and di-liganded (right) nAChR structures, with the dashed red line indicating in each case the release of backbone conformational restraints.

The di-liganded ACh-bound structure also suggests the molecular basis for the slightly increased affinity and efficacy of ACh relative to carbamylcholine^25,26^. Although both ACh and carbamylcholine bind to the nAChR with similar poses and induce a similar collapse of the aromatic box, the movement of loop F from the γ subunit is larger in the ACh-bound structure, positioning γAsp174 and γGlu176 to interact more strongly with loop C αThr191 and β7 αLys145, respectively (Extended Data Fig. 8). These enhanced interactions likely stabilize the agonist-bound activated state.

### The mono-liganded nAChR with α_γ_ activated and α_δ_ primed

The mono-liganded structure, with ACh bound to the α_γ_γ but not the α_δ_δ agonist site, exhibits features both characteristic of and intermediate to the unliganded and di-liganded structures (Figs. 2-4). At the level of the agonist sites, ACh binding to the α_γ_γ agonist site stabilizes an activated-like conformation where the degree of loop C capping around the bound ACh is close to that observed in the di-liganded structure. The distance between the Cα carbons of α_γ_Cys192 and γGlu56, residues located on either side of the binding site, decreases from 17.6 Å in the unliganded structure to 10.0 Å and 9.1 Å in the mono-liganded and di-liganded structures, respectively (dashed line in Fig. 2a, zoomed in view). There is a similar contraction of the aromatic box around the quaternary amine of ACh, with the area of the quadrilateral formed by the Cα carbons of αTrp149, αTyr190, αTyr198, and γTrp55 decreasing from 109 Å^2^ in the unliganded structure to 79 Å^2^ and 73 Å^2^ in the mono-liganded and di-liganded structures, respectively (Fig. 2c). Loop F from the complementary γ subunit moves substantially towards its capped conformation in the di-liganded structure, although there are limited interactions between γAsp174/γGlu176 (loop F) and αThr191/αLys145 (loop C/β7) (Fig. S7). With loop C and the four aromatic box residues essentially trapping the bound ACh, the α_γ_γ agonist site adopts an activated high affinity ACh-binding conformation.

In contrast, loop C at the α_δ_-δ agonist site remains uncapped, with the distance between the Cα carbons of α_δ_Cys192 and δAsp59 only decreasing from 18.0 Å in the unliganded structure to 17.4 Å versus 8.7 Å in the mono-liganded and di-liganded structures, respectively (Figs. 2 and Extended Data Fig. 7). There is essentially no contraction of the α_δ_δ aromatic box, with the quadrilateral formed by the Cα carbons of αTrp149, αTyr190, αTyr198, and δTrp57 only decreasing from 107 Å^2^ in the unliganded state to 106 Å^2^ versus 75 Å^2^ in the mono-versus di-liganded states, respectively (Fig. 2c). Loop F from the complementary δ subunit only moves slightly from its uncapped position. Despite ACh binding to the α_γ_γ agonist site, the α_δ_δ agonist site remains in a solvent accessible resting-state-like low affinity conformation. The α_γ_γ and α_δ_δ agonist sites are structurally ‘independent’, consistent with the lack of cooperativity in ACh binding to the *Torpedo* nAChR suggested by a rectangular hyperbolic equilibrium binding curve^16^.

At the level of the interface between the extracellular and transmembrane domains, the α_γ_-subunit adopts a locally active conformation while α_δ_ adopts a conformation along its trajectory towards the Leu9’ pore open state. Specifically, agonist-binding to the α_γ_γ agonist site correlates with a pivoting of the entire α_γ_ extracellular domain (Movie 1) so that α_γ_Pro265 (M2-M3 loop) translates outward from the pore-proximal side to the pore-distal side of α_γ_Val46 (β1-β2 loop) to occupy a position equivalent to that observed in the di-liganded structure (Fig. 3)^19^. On the other hand, even though the α_δ_δ agonist site is unoccupied and the α_δ_ extracellular domain remains essentially in a resting state-like conformation, α_δ_Pro265 is positioned below α_δ_Val46, a position between its pore-proximal and pore-distal positions in the unliganded and di-liganded structures, respectively. The outward movement of α_δ_Val46, and thus of the α_δ_ M2-M3 loop, along its trajectory towards the di-liganded structure suggests that despite the unoccupied α_δ_δ agonist site, the interface between the extracellular and transmembrane domains of α_δ_ is poised for activation.

Note that the relative positions of the αVal46/αPro265-equivalent residues in non-α subunits in the mono-liganded structure are all indicative of a locally ‘active’ conformation in that each αPro265-equivalent residue is positioned on the pore distal side of the αVal46-equivalent residue (Extended Data Fig. 9). This locally active conformation, however, is observed in all non-principal agonist binding subunits in both unliganded and fully liganded structures of the nAChR and other pLGICs^19^. As noted previously, agonist binding to heteromeric pLGICs leads to a relaxation of local asymmetry at the inter-domain interface to a more symmetric conformation^19^.

At the level of the channel pore, the α_γ_-subunit adopts a Leu9’ gate open conformation while the transmembrane domains of the other subunits, including α_δ_, adopt conformations intermediate between their conformations in the closed resting and di-liganded channel Leu9’ gate open states. Specifically, activation of the α_γ_γ agonist site correlates with the same outward twisting/tilting of the pore-lining α_γ_M2 α-helix that is observed in the di-liganded structure, with key hydrophobic α_γ_M2 residues, such as the α_γ_Leu9’ gate, adopting equivalent positions in both mono-liganded and di-liganded structures (Fig. 4 and Movie 2). Significantly, activation of the α_γ_ transmembrane domain correlates with movements of other pore-lining M2 α-helices, each adopting a position along its trajectory leading to the di-liganded pore open state. Even the C-terminal half of the α_δ_ M2 α-helix tilts/twists away from the pore, although the rotameric position of its key Leu 9’ gate residue still superimposes on its position in the unliganded structure. The mono-liganded structure has a de-wetted pore that does not conduct cations across the membrane (Fig. 4 and Extended Data Fig. 10). Interestingly, detectable water penetrations are frequently observed in the mono-liganded structure prior to the release of conformational restraints.

In summary, agonist binding to the α_γ_γ agonist site in the mono-liganded structure stabilizes a conformation where the entire α_γ_-subunit fully transitions to an activated conformation, while the unoccupied α_δ_ principal and other subunits remain inactive, albeit poised for activation. The mono-liganded structure corresponds to a mono-liganded ACh-bound pre-open intermediate state.

### A structure-guided scheme reconciles single-channel and structural data

Modern kinetic schemes describing nAChR single-channel activity predict the existence of pre-open intermediate states, with the rate constants inferred from maximum-likelihood fits of single channel recordings^5,27^ making it possible to calculate the steady-state probabilities of pre-open intermediates and thus the ACh concentrations at which they should prevail. To extend our findings to the human nAChR and assess whether current schemes correctly predict the prevalence of the mono-liganded pre-open intermediate observed by cryo-EM, we recorded single-channel data at ACh concentrations spanning both the ‘conformational response’ and the observed macroscopic activation of the human adult muscle-type nAChR (Fig. 5). We globally fit bursts of single-channel activity to two established kinetic schemes, the ‘flip’^3^ and ‘prime’^5^ models, each of which explicitly incorporates a mono-liganded pre-open intermediate (Extended Data Figs. 11 and 12). Attempts to fit the data to the simpler ‘flip’ model, which stipulates that the two agonist sites in the resting state have identical ACh association and dissociation rate constants (i.e., identical ACh binding affinities)^27^, failed to converge to a unique set of values. Fitting to the ‘prime’ model, where the agonist site affinity constraints are relaxed, converged to a set of rate constants similar to those published previously (Extended Data Fig. 11 and Table 3). The inferred rate constants, however, predict that amongst the nAChR conformational ensemble, the steady-state proportions of receptors in the mono-liganded primed state are vanishingly small at all concentrations across the ACh-induced response (Extended Data Fig. 12). Given that the macroscopic and single-channel behaviour of the *Torpedo* and human nAChR are similar (Figs. 1a, 5a, and Extended Data Fig. 14), the predicted steady state proportions of the mono-liganded primed state indicate that it would be difficult, if not impossible, to resolve this state by cryo-EM, at any ACh concentration. In its present form, the prime model is incompatible with our observation that the mono-liganded pre-open state represents 5-10% of the total particles imaged at sub-saturating ACh concentrations (Figs. 1 and Extended Data Fig. 5).

**Figure 5.**
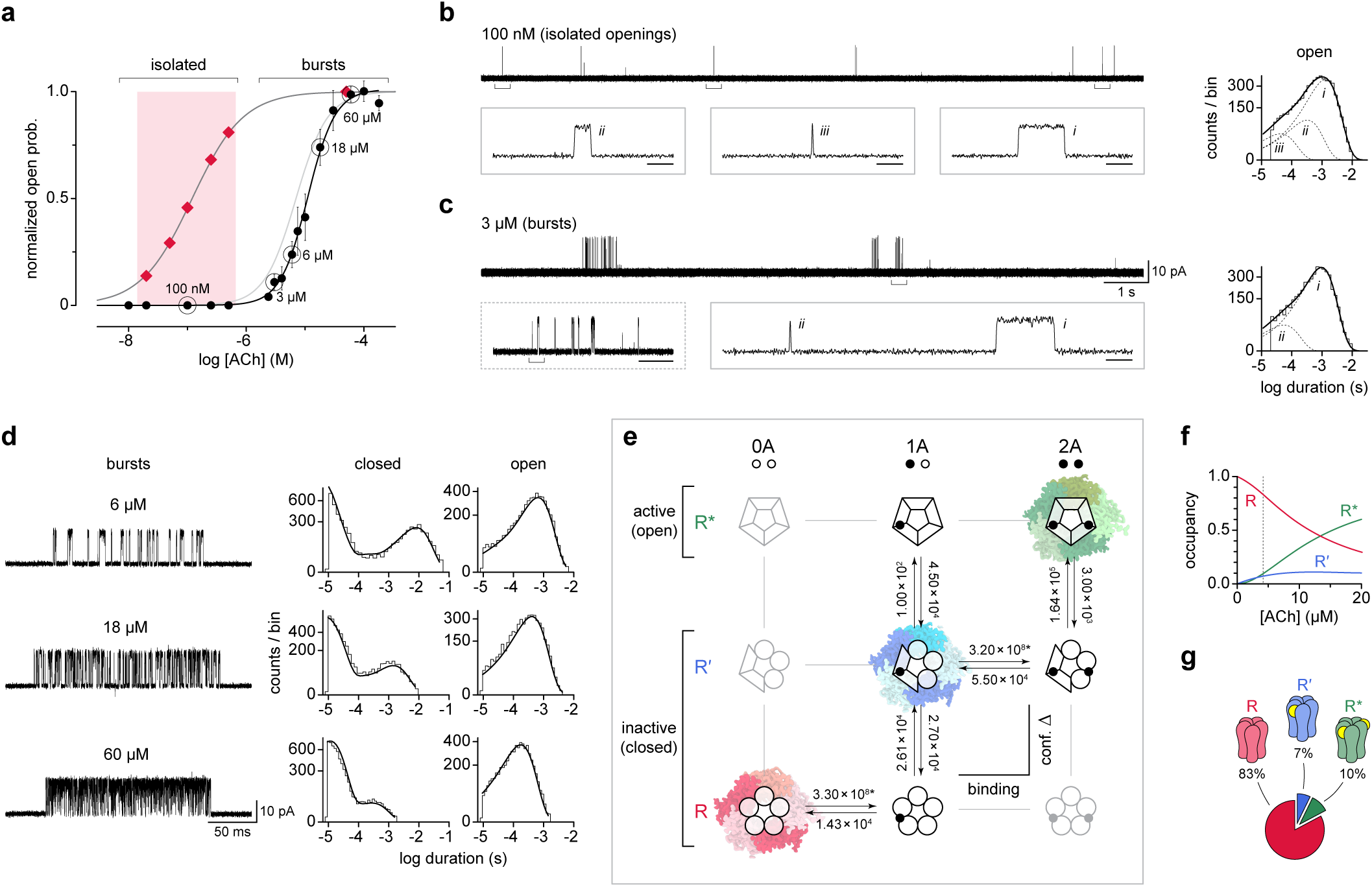
Single-channel activity of the human muscle-type acetylcholine receptor predicts the steady-state occupancy of the mono-liganded intermediate state observed by cryo-EM. **(a)** Normalized open probability of bursts of single-channel activity from the human adult muscle-type acetylcholine receptor (black). Data points represent mean burst open probability with error bars being one standard deviation from the mean (see Table S4 for statistics). Single-channel activity for the indicated (circles) acetylcholine concentrations are shown in subsequent panels. Also, shown are the fits of the ethidium bromide (dark gray) and human whole-cell (light gray) dose responses from Figure 1, as well as the acetylcholine concentration range (pink box) at which the cryo-EM data sets were collected (red diamonds). **(b, c)** Single-channel activity from cell-attached patches in the presence of **(b)** 100 nM acetylcholine occurs as individual openings in relative isolation, whereas at **(c)** 3 µM acetylcholine or above, successive openings coalesce into bursts. The scale bar in ‘c’ applies to the continuous traces in both panels. Insets are expansions of the bracketed regions and the depicted openings (*i,ii,iii*) reflect the mean durations of the components (dashed lines) in the associated open duration histograms (right). The height of the inset boxes represents 30 pA in each case, where the horizontal scale bar represents 50 ms (dashed box in ‘c’) or 1 ms (solid boxes in ‘b’ and ‘c’). **(d)** Single-channel bursts of human adult acetylcholine receptor activity (left) with corresponding open and closed duration histograms (right) for select acetylcholine concentrations. Global fits (solid lines; see also Figure S11) to the scheme presented in ‘e’ overlay the histograms in each case. **(e)** Structure-informed ‘sequential’ scheme used to describe the single-channel activity. Theoretical states not included in kinetic fitting are faded, while cryo-EM densities of touchstone states solved in the present work are also shown. Inferred rate constants for each transition (see Table S3 for units and error estimates) are used to predict **(f)** the acetylcholine concentration dependent state occupancy of receptors in a resting (R; red), intermediate (Ŕ; blue), or active (R*; green) conformation. **(g)** At low acetylcholine concentrations (dashed line in ‘f’), the inferred rates predict the steady state occupancies observed in the cryo-EM data set. All single-channel data was recorded at –120 mV and presented filtered at 10 kHz. Detections for kinetic analysis were made at 25 kHz (see Figures S11 & S13), where openings are upward deflections.

As emphasized by Colquhoun, kinetic models used to fit single-channel data “*are useful only in so far they are a sufficiently good approximation of the actual physical reality of the receptor’s conformations*”^3,28^. Our structures show that ACh binding to one agonist site leads to an intermediate conformation where the nAChR is partially activated prior to ACh binding to the second agonist site. To reconcile this structural observation with our single-channel data, we simplified the ‘prime’ model by removing the di-liganded resting state – a simplification that imposes the ‘bind-prime-bind’ trajectory elucidated by the structural data. Global fitting of the entire single-channel data set to this simplified, but structure-guided scheme, robustly fits the observed single channel activity (Extended Data Tables 2 and 5). Notably, without imposing any constraints, the association rates for ACh binding to the two agonist sites converge to virtually identical values, while the dissociation rates differ by less than four-fold, both sets of values in line with the observed structures and conformations of the two agonist sites (Extended Data Table 3)^5,29^. In particular, the near identical association rate constants are consistent with the two sites being uncapped and thus equally accessible to agonist prior to binding. In addition, the inferred rate constants predict that a mono-liganded intermediate state represents as much as 10% of the nAChR conformation ensemble at sub-saturating ACh concentrations (Figs. 5f, 5g and Extended Data Fig. 12). Despite the inherent limitations of extrapolating these predictions to the nanomolar ACh concentrations used to collect our cryo-EM data sets, particularly given that the cryo-EM data were recorded using the *Torpedo* nAChR reconstituted into MSP2N2 nanodiscs, as opposed to the human adult nAChR expressed on the surface of cells (see Methods), the prediction that close to 10% of the nAChRs adopt a mono-liganded intermediate state at sub-saturating ACh concentrations aligns remarkably well with the proportion of mono-liganded particles observed by cryo-EM (Fig. 1a, 5g). Thus, the sequential ‘bind-prime-bind’ trajectory revealed by our structural data not only informs the kinetic scheme used to describe the single-channel data, it also leads to a simple, unifying model that robustly describes nAChR activation.

## Discussion

We have combined single-molecule structural data with single-channel electrophysiological recordings to provide new structural and mechanistic insight into acetylcholine receptor activation. We solved three structures of the muscle-type nAChR from *Torpedo* along the pathway leading from the unliganded closed, to a di-liganded activated state. Importantly, we isolated a structure with ACh bound to only one of the two agonist-binding sites. This structure shows that ACh binding exclusively to the α_γ_γ agonist site stabilizes a state where the α_γ_-subunit has transitioned to an activated conformation, with α_γ_ activation leading to a restructuring of the entire transmembrane domain to an intermediate conformation that appears poised for activation. Additional binding of ACh to the α_δ_δ agonist site is required to stabilize a conformation where α_δ_, along with the other subunits, are fully activated leading to hydration of the ion channel pore. Based on these structural observations we introduce a new mechanistic scheme for activation that both describes the single-channel activity of the human nAChR, and accurately predicts the steady-state probabilities of unliganded, mono-liganded, and di-liganded receptors that matches the proportion of particles observed in each state in the cryo-EM maps. Together, the structural and functional data delineate an activation mechanism where ACh binding and individual subunit conformational changes are both sequential and asynchronous.

The key aspect of the present work is the direct observation by cryo-EM of a mono-liganded intermediate pre-open state. Intermediate pre-open states were first detected in single-channel recordings and employed to improve kinetic fitting of related glycine receptors^7,30^, and subsequently the muscle-type nAChR^3,31^. In the first scheme to incorporate an intermediate state, the ‘flip’ model, the receptor transitions from the resting state to an intermediate ‘flipped’ state before it can open. One postulate of the flip model is that the conformational change of subunits is concerted, with all subunits simultaneously adopting either a resting, ‘flipped’, or open conformation^7^. This postulate was relaxed in the subsequent ‘prime’ model, which proposes that individual nAChR subunits, or its two agonist sites, ‘prime’ independently^4^. Further refinements converged upon a ‘reduced prime model’ with only a single/initial level of ‘priming’^5^. Building off this reduced prime model, the scheme developed here preserves the independent conformational changes of subunits but does not include a di-liganded resting state. Within this new scheme, the mono-liganded pre-open state detected by cryo-EM represents a mono-liganded ‘primed’ conformation. Indeed, the observation that loop C of the α_γ_γ agonist site is fully capped is consistent with the original disulfide trapping experiments^4^. Furthermore, ACh occupancy of the α_γ_γ site not only stabilizes the α_γ_-subunit in an activated conformation, but also correlate with subunit-specific conformational changes that have propagated to much of the remaining transmembrane domain leading to a channel that, although closed, is clearly poised (or ‘primed’) for full activation. For these reasons, we propose that the mono-liganded intermediate pre-open structure resolved by cryo-EM represents a mono-liganded ‘primed’ state.

If indeed a primed state, the mono-liganded structure sheds light on the structural changes associated with priming. Intermediate structures between apo and activated states have been solved for homomeric pLGICs^8–12^. Although not attributable to primed states, these structures all exhibit an extracellular domain that has transitioned to an active or active-like conformation, while the transmembrane domain remains predominantly inactive. These structures suggest that priming occurs through a concerted mechanism, whereby entire domains activate sequentially (i.e., the entire extracellular domain activates prior to the entire transmembrane domain). In stark contrast, the mono-liganded structure presented here shows that the heteromeric nAChR exhibits asynchronous transitions of individual subunits, as opposed to entire domains. Whether this asynchronous conformational change of subunits is unique to heteromeric pLGICs remains to be seen. New structures of homomeric pLGICs in an intermediate state with low agonist occupancy are required.

Only one mono-liganded structure, with ACh bound exclusively to the α_γ_γ agonist site, was detected in the cryo-EM data sets. On the surface, this observation is consistent with the α_γ_γ site having a higher apparent affinity for ACh than the α_δ_δ site^32–34^, an interpretation in line with the noted ensemble of loop C capping interactions at α_γ_γ, which are absent or weaker at α_δ_δ^23,35^. Parsimoniously, our scheme describing bursts of adult human nAChR single-channel activity only includes one mono-liganded primed, and one mono-liganded open state. By itself the scheme does not specify which of the two agonist sites is occupied in these mono-liganded states. Given the mono-liganded cryo-EM structure, however, the simplest interpretation is that ACh is bound to the α_γ_γ site (or α_ε_ε in the adult human nAChR) in the mono-liganded primed state. If correct, this leads to a mechanism where binding and conformational change of individual subunits appears to be sequential and initiated through ACh binding first to the α_γ_γ (or α_ε_ε) site.

It should be noted however, that although the presented scheme reconciles the available data, it is undoubtedly an oversimplification of a comprehensive framework that incorporates additional states and transitions. For example, unliganded priming and activation are almost certainly possible (as shown in Fig. 5e), as is an additional di-liganded priming step akin to the one observed in the original prime model. Furthermore, binding to the α_δ_δ agonist site first, is also possible. Indeed, at ACh concentrations below the threshold for initiating bursts, it is well-established that two classes of isolated mono-liganded openings, each thought to originate from single occupancy of one or the other agonist site, are observed (Fig. 5b)^36,37^. The probability of observing a particular state depends not only on the states lifetime, but also the individual rates along the path to that state. Our inability to detect α_δ_δ mono-liganded states by cryo-EM, or at bursting concentrations in single-channel recordings, may simply relate to relative rates of binding/priming/activation from the α_γ_γ (α_ε_ε) versus α_δ_δ site. Regardless of the exact mechanism, the important point is that while these additional states and transitions are theoretically possible, their inclusion in our scheme is not necessary to explain our observations nor are they likely required to describe the wild type post-synaptic response. Moving forward it will be interesting to see if such additional states can be exposed through mutation, or the use of partial agonists.

The mono-liganded structure helps explain the paradoxical observation that nAChR activation appears cooperative (Hill coefficient = 1.78 ± 0.35, n=87) while ACh binding does not (see Results^16^). In the context of activation, the mono-liganded structure shows that ACh binding to α_γ_γ not only leads to activation of the entire α_γ_-subunit, but also additional conformational changes in the remaining subunits, which shifts them towards their active conformation, despite the pore remaining closed. Although the transmembrane domain is clearly poised for full activation, it takes binding of a second ACh molecule to fully drive the conformational transition, and thus the two agonist sites work together to stabilize the fully activated/open state, accounting for the observed cooperativity of channel activation. On the other hand, in the context of ACh binding, the mono-liganded structure shows that ACh binding to α_γ_γ has little effect on the structure of the α_δ_δ site. Specifically, ACh binding to the α_γ_γ site leads to minimal capping of loops C and F in α_δ_δ and thus little contraction of the α_δ_δ site. From a structural perspective, the conformations of the α_γ_γ and α_δ_δ agonist sites appear independent of one another, likely accounting for the apparent lack of cooperativity in ACh binding.

Finally, the structures presented here shed light on the global nature of allosteric transitions in the muscle-type nAChR. Activation has often been interpreted in terms of a MWC-like allosteric model^38^, which posits that the energies between the inactive/closed and active/open states govern their relative proportions, and thus the probability of the channel being in the open state^39^. In the absence of ACh, the resting state has a lower energy and thus predominates with only rare excursions to the open state. Energy derived from preferential binding of either one or two molecules of ACh, stabilizes the open conformation, with this energetic stabilization being greater in the di-liganded than in the mono-liganded state. Although these energetic tenets of the MWC model are demonstrably applicable to the nAChR in the context of the two end states, closed and open, a central tenet of the MWC model is that conformational symmetry of subunits is maintained, and thus that the subunits in oligomeric proteins, such as the nAChR, undergo concerted all-or-none conformational transitions^40^. The finding that ACh occupancy in the α_γ_γ site stabilizes a conformation where the entire α_γ_-subunit has transitioned into an activated-like conformation, while much of the α_δ_ subunit remains in a resting conformation, demonstrates that conformational transitions in the heteromeric *Torpedo* nAChR are not ‘all or none’, nor do they occur via a concerted and symmetric transition. Instead, asynchronous conformational transitions can be demonstrably observed and quantified.

---

**Movie 1. Top-down view of the conformational transition in the extracellular domain from the unliganded to the mono-liganded, and then to the di-liganded state.** A global alignment of the three states was done based on the C_α_ atoms in the M1-M3 helices from each subunit to visualize the motions of the extracellular domain relative to the transmembrane domain. A morph between the three states was created in ChimeraX^41^.

**Movie 2. Top-down view of the conformational transition in the transmembrane domain from the unliganded to the mono-liganded, and then to the di-liganded state.** A global alignment of the three states was done based on the C_α_ atoms within the β-strands from each subunit to visualize the motions of the transmembrane domain relative to the extracellular domain. A morph between the three states was then created in ChimeraX^41^. Tan sticks in the channel pore correspond to the Leu9’ and Val13’ residues from each subunit that form the hydrophobic gate.

## METHODS

### Protein purification and reconstitution into nanodiscs

The nAChR was purified from the electroplax tissue (Aquatic Research Consultants, CA) of the Pacific electric ray *Tetronarce californica* (NCBI: txid7787) using a bromoacetylcholine bromide-derivatized Affi-Gel 102 column (Bio-Rad) as previously described^1^, but with the following modifications. After the nAChR was eluted from the column, it was incubated with 15 mM DTT to break disulfide-linked pentamers prior to purification on a Superose6 size-exclusion column. The purified monomeric form of the nAChR pentamers was incubated with soybean azolectin and MSP2N2^2^ and then dialyzed (12-14 kDa cut-off) five times against 2 L of tris dialysis buffer, with buffer changed roughly every 12 h. The MSP2N2 nanodisc-reconstituted nAChR was further purified by size-exclusion chromatography, with the appropriate fractions pooled, concentrated, aliquoted, snap frozen in liquid nitrogen and stored at −80°C. To ensure removal of any residual agonist remaining from the nAChR affinity column elution, aliquots were diluted 75-fold in tris dialysis buffer, incubated for 30 minutes on ice with occasional mixing, and concentrated back to the original volume, with the process repeated five times. The buffer exchanged nAChR was injected once again on the size-exclusion column and the monomeric nAChR peak pooled. The pooled fractions were again concentrated to 0.65 mg/mL and snap frozen.

### Fluorescence measurements of ACh-activation

Fluorescence experiments were performed essentially as described previously^3^ on a Cary Eclipse fluorescence spectrophotometer (Varian Inc.) with the ethidium fluorescence excited at 500 nm (±5 nm slits) and the fluorescence emission intensity measured at 590 nm (±20 nm slits). Increasing concentrations of ACh were added every 2 minutes to cuvettes containing 0.1 μM EthBr either with or without the nanodisc-reconstituted nAChR. The fluorescence emission was recorded at each concentration of ACh. Each plot of the increase in fluorescence versus concentration of ACh was fit with a variable slope sigmoidal dose-response curve to determine the EC_50_ of the ACh-induced conformational response in GraphPad Prism, version 9.

### Electron microscopy and image analysis

The nAChR in asolectin-MSP2N2 nanodiscs was mixed with the megabody Mbc7HopQNbF3^4^, which binds MSP2N2 not the nAChR, at a molar ratio 1:3 to overcome orientation bias of the nAChR on the cryo-EM grids. After incubating with the desired concentration of ACh for 30 min on ice, 3.5 μls of each sample were deposited onto a glow-discharged (30 mA, 50 s) Quantifoil Au/C R 1.2/1.3 grid, blotted for 6 s with force 0 at 8 °C and 100% humidity, and then plunge-frozen in liquid ethane using a Mark IV Vitrobot (Thermo Fisher Scientific).

All cryo-EM datasets were recorded on a Glacios electron microscope at the Institut de Biologie Structural in Grenoble, France. Movies (40 frames) were recorded at a nominal magnification of 36,000 with SerialEM and a Gatan K2 Summit camera. The raw movies were imported to Cryosparc^5^, aligned, and summed. CTF estimation was calculated for the non-dose-weighted sums. Particles were autopicked with crYOLO 1.7.6^6^ using the general model for low-pass filtered images. Particle coordinates were imported into Cryosparc, where all subsequent data processing was performed. Raw particle stacks were extracted using 256 pixel box sizes.

Each dataset was initially subjected to two rounds of 2D classifications, *ab initio* reconstruction, and heterogeneous refinement (3 classes). The class showing the best nAChR features (map integrity and isotropic viewing angle distributions) was selected and refined using non-uniform and local refinement^7^ to yield a consensus reconstruction. Each consensus reconstruction was further assessed for conformational heterogeneity using 3D classification (10 classes) and 3D variability analysis (3DVA) with intermediates (30 frames) and clusters (8 clusters)^8^, in each case using a single focused mask that encompasses regions of the nAChR that change most between apo and agonist bound states (the two agonist-binding sites, all five pore-lining M2 α-helices, and the α_δ_ M4 α-helix).

To obtain a structure of the mono-liganded state, particles from the consensus refinements of all five data sets recorded at sub-saturating concentrations of ACh (2,461,091 particles) were combined and subjected to 3D classifications and 3DVA, with the particles sorted into each of the ten classes/eight clusters refined as described above. Particles contributing to clusters/classes that did not clearly resemble either un-liganded pore closed or di-liganded pore open states (1,494,365 particles) were re-combined and subjected to a second round of 3D classifications and 3DVA. After the second round, the remaining 772,485 non-end state particles (i.e., clearly not un-liganded or di-liganded) were again refined, with a subsequent 3D classification and 3DVA cluster analysis still yielding reconstructions corresponding to un-liganded pore closed, di-liganded pore-open states, and a mono-liganded state with a capped α_γ_ loop C and an uncapped α_δ_ loop C.

To remove remaining particles corresponding to the two end states, the best refined maps from the 3D classifications for the un-liganded, mono-liganded, and di-liganded states were selected and used as inputs for another round of 3D classification and heterogeneous refinement. 379,957 particles were sorted into the mono-liganded state. The resulting map generated from these particles was refined and subjected to a second round of 3D classifications and 3DVA. The corresponding density maps revealed a mono-liganded state with some remaining heterogeneity in the α_γ_ loop C. To further ensure all particles in the reconstruction corresponded to the mono-liganded state, a third round of 3D classification with the same input maps was performed and the remaining 129,828 particles sorted to the mono-liganded state. The resulting map generated from these particles was refined to obtain a well-defined 3.13 Å resolution reconstruction of the mono-liganded state.

### Model refinement and structure analysis

Preliminary model building was performed in Isolde^9^. Cycles of real-space refinement were then performed in Phenix^10^ with manual rebuilding in Coot^11^ using the unsharpened maps from Cryosparc. Occasionally, maps post-processed with DeepEMhancer^12^ were used during manual building in Coot to aid in modelling poorly resolved side chains. Validation was performed using the unsharpened maps with Molprobity^13^.

### Two-electrode voltage clamp electrophysiology

Oocytes from *Xenopus laevis* were injected with *Torpedo*/human adult wild type subunit cRNA as described previously^1^ and allowed to incubate at 16 °C in ND96+ buffer (5 mM HEPES, 96mM NaCl, 2 mM KCl, 1 mM MgCl_2_, 1 mM CaCl_2_, 2 mM pyruvate). Whole-cell currents were measured in response to agonist concentration jumps (flow rate of 5-10 mL/min.) using a two-electrode voltage clamp apparatus (OC-725C oocyte clamp; Holliston, MA) in the presence of 1 mM atropine. The whole-cell currents were recorded in HEPES buffer (96 mM NaCl, 2 mM KCl, 1.8 mM BaCl_2_,1 mM MgCl_2_, and 10 mM HEPES, pH 7.3), with the transmembrane voltage clamped at −60 mV.

### Mammalian cell expression

The full complement of human adult muscle-type nAChR subunit cDNAs (α1, β1, δ, and ε) were transfected into BOSC 23 cells^14^, originally from ATCC (CRL11270), but provided by Steven M. Sine (Mayo Clinic) (RRID:<u>CVCL_4401</u>). Cells were maintained in Dulbecco’s modified Eagle’s medium (DMEM; Corning) containing 10% (vol/vol) fetal bovine serum (Gibco) at 37 °C, until they reached 50–70% confluency. Cells were then transfected using calcium phosphate precipitation, and transfections terminated after 3–4 h by exchanging the medium. All experiments were performed within one day post transfection (i.e., within 24h after exchanging the medium). A separate plasmid encoding green fluorescent protein was included in all transfections to facilitate identification of transfected cells. For *Torpedo* nAChRs, cDNAs encoding the α1, β1, δ and γ-subunits of *T. marmorata* were provided by Steven M. Sine (Mayo Clinic). Cells were maintained and transfected as described above, except that 12 h post transfection with *Torpedo* cDNAs, cells were transferred to 26 °C and allowed to express for 48 h before experiments were conducted.

### Single-channel patch clamp recordings

Single-channel recordings from BOSC 23 cells transiently transfected with cDNAs encoding wild-type subunits, were obtained in the cell-attached configuration with a membrane potential of –120 mV and a temperature maintained between 19 and 22 °C. The bath solution contained (in mM) 142 KCl, 5.4 NaCl, 0.2 CaCl_2_ and 10 HEPES (4-(2-hydroxyethyl)−1-piperazineethanesulfonic acid), adjusted to pH 7.40 with KOH. The pipette solution contained (in mM) 80 KF, 20 KCl, 40 K•aspartate, 2 MgCl_2_, 1 EGTA (ethylene glycol-bis(β-aminoethyl ether)-*N*,*N*,*N*′,*N*′-tetraacetic acid), and 10 HEPES, adjusted to a pH of 7.40 with KOH. Acetylcholine chloride (Sigma) was added to pipette solutions to the desired final concentration and stored at –80 °C. Patch pipettes were fabricated from type 7052 or 8250 nonfilamented glass (King Precision Glass) with inner and outer diameters of 1.15 and 1.65 mm, respectively, and coated with SYLGARD 184 (Dow Corning). Prior to recording, electrodes were heat polished to yield a resistance of 5 to 8 MΩ. Single-channel currents were recorded using an Axopatch 200B patch clamp amplifier (Molecular Devices), with a gain of 100 mV/pA and an internal Bessel filter at 100 kHz. Data were sampled at 1.0 μs intervals using a BNC-2090 A/D converter with a National Instruments PCI 6111e acquisition card and recorded by the program Acquire (v. 6.0.0; Bruxton).

### Cell line authentication and mycoplasma testing

Approximately five million confluent cells were harvested and their total DNA isolated (E.Z.N.A.® Tissue DNA Kit), and then submitted to The Centre for Applied Genomics Genetic Analysis Facility (The Hospital for Sick Children, Toronto, Canada) for STR profiling using Promega’s GenePrint® 24 System. A similarity search on the 8159 human cell lines with STR profiles in Cellosaurus release 42.0 was conducted on the resulting STR profile, which revealed that the cell line shares closest identity (88%, CLASTR 1.4.4 STR Similarity Search Tool score) with Anjou 65 (CVCL_3645). Anjou 65 is a child of CVCL_1926 (HEK293T/17) and is itself a parent line of CVCL_X852 (Bartlett 96). Bartlett 96 is the parent line of BOSC 23^14^. PCR tests confirmed that the cells were free from detectable mycoplasma contamination.

### Single-channel analysis and kinetics

To measure the step response of our Axopatch 200B patch clamp amplifier, a Tektronix AFG1062 arbitrary function generator was connected to the Speed test input of the amplifier, and rectangular pulses with a 40 mV amplitude at 1 kHz frequency were recorded to the hard disk of a PC computer while sampling at 1 µs intervals using the program Acquire (v. 6.0.0; Bruxton). 10,000 individual pulses were aligned and averaged within R^15^, to obtain the mean step response with an improved signal-to-noise ratio. This averaged step response was then fit with a sigmoidal dose response curve (variable slope) within GraphPad Prism V.8.4.3, and the 10% to 90% risetime calculated (14.513 µs at 25 kHz) to provide and accurate measure of the instrument dead time (7.808 µs) (dead time = risetime x 0.538)^16^.

Single-channel event detection was performed using the program TAC (v. 4.3.3; Bruxton), where recordings were analysed with an applied 25 kHz digital Gaussian filter. Opening and closing transitions were detected using the 50% threshold crossing criterion. Histograms were visually fit with a minimum sum of exponential components. Closed duration histograms were used to calculate a critical closed duration (τ_crit_), defined as the intersection of the slowest activation and the fastest desensitization components^17,18^. Closings beyond the critical closed time were discarded. Events were imported into R using *scbursts* (v. 1.6)^19^, where individual durations were corrected for the instrument risetime, and then individual bursts defined by τ_crit_. Bursts with less than three openings were removed from downstream analysis. The open channel probability (pOpen) was calculated for each burst, and bursts with a pOpen that did not fit within a normal distribution were removed with *extremevalues* (v. 2.3.3)^20^. The resulting bursts were then fit with a single Gaussian based on their pOpen, and bursts beyond 2 standard deviations from the mean were removed from further kinetic analysis.

Kinetic schemes were fit to the sequence of single-channel dwells in the global dataset using maximum likelihood fitting implemented within MIL (QUB suite, State University of New York, Buffalo, NY), with a user-defined dead time of 7.808 μs. MIL corrected for missed events, estimated and gave standard errors of all model parameters by maximum likelihood. Histograms were generated from resulting fits and exported from R using *scbursts* (v 1.6). Each kinetic scheme was fit to the sequence of dwells within MIL, which provided a log likelihood (LL) value describing how well the model fit the data. Models were compared using a combination of log likelihood ratio tests and the Schwarz Informational Criterion (LLRs, SICs; Table S5)^21,22^. LLRs were calculated as LLR = LL*_k1_* – LL*_k2_*, where ‘*k_1_*’ corresponds to the model with more free parameters (i.e., is more complex) and ‘k_2_’ is the model with fewer free parameters (i.e., the simpler model). Significance was determined by comparing twice the LLR statistic to the χ^2^ distribution as described previously (ref. 57 and 58). Schwarz Information Criterion values (SICs) were calculated to rank different models, with the smallest number ranking the highest^23^. The SIC for each model was calculated from SIC = -LL + (0.5*F*)*ln(*N*), where ‘*F*’ is the number of free parameters, and ‘*N*’ is the number of observed events (e.g., openings and closings). Both LLR and SIC tests provide statistical support for the reduced prime model (Table S5), however as the inferred rates predict negligible steady-state occupancy of the mono-liganded primed state, the reduced prime model is incompatible with the cryo-EM data. Conversely, the sequential model, which is a simplified subset of the reduced prime model, [1] still explains the single-channel activity, [2] correctly predicts the steady-state occupancy of the mono-liganded primed state, and [3] leads to inferred agonist association rates that are compatible with the observed structure and conformations of the two uncapped agonist sites.

Event detection for low ACh concentrations (100 nM ACh; Figure 5) were analysed with an applied 10 kHz digital Gaussian filter. Data was pooled from several recordings until ∼5,000 openings were collected. Open duration histograms were fit by exponential components within TACFit. LLR’s and SICs were performed to validate the inclusion of additional exponential components (Table S2)^23^.

State probability plots were generated using the QUB Mandelics Suite^24^ by inputting a user defined kinetic scheme with specified rate constants. Steady-state state probabilities were calculated across the entire concentration range, and the results plotted in GraphPad Prism V.8.4.3 and paths 4% smoothed in Adobe Illustrator v.29.0.1.

### MD Simulations

Glycosylation chains and lipids were removed from the structures of the unliganded, mono-liganded and di-liganded states used as simulation inputs, with ACh kept in the binding sites for the mono-liganded and di-liganded states. Systems were set up using CHARMM-GUI membrane builder^25^. Each structure was embedded in a 140×140 Å bilayer containing a 3:1:1 ratio of POPC:POPA:Cholesterol, with a box length of 190 Å. Disulfide bonds were preserved.

Simulations were performed using the NAMD 3 simulation engine^26^ with the CHARMM36 forcefield^27,28^ applied to the proteins, ions, lipids and ACh using the TIP3P^29^ water model and a NaCl concentration of 150mM. Each system was minimized for 1000 steps. Systems were equilibrated for 5 ns with soft harmonic positional restraints on all heavy protein atoms, followed by 50 ns of equilibration with positional restraints on heavy backbone atoms only. The temperature was held at 300 K using the Langevin thermostat^30^ and the pressure held at 1 atm using the Langevin piston method. Covalent bonds, including hydrogen atoms, were constrained using the SHAKE/RATLLE algorithms^31,32^. The SETTLE algorithm^33^ was used for water. Long-range electrostatic interactions were evaluated by the particle-mesh Ewald (PME) algorithm^34^.

Production runs were performed with no positional restrains for five repeats of 200 ns for each system. Hydrogen mass repartitioning^35^ was used for all production runs allowing for a timestep of 4 fs. A timestep of 8 and 4 fs was used for long- and short-range interactions, respectively, using the r-RESPA multiple time-stepping algorithm^36^.

The MDAnalysis python package^37,38^ was used to create in-house scripts for simulation analysis. Pore diameter was measured using the HOLE program^39^. Structural figures were prepared using VMD^40^ and ChimeraX^41^. Plots were prepared using the python package matplotlib^42^.

## Data Availability

All atomic models and cryo-EM maps have been deposited in the Protein Data Bank and Electron Microscopy Data Bank: unliganded (PDB 9E3F, EMDB 47481); mono-liganded (PDB 9E3G, EMDB 47482); and di-liganded (PDB 9E3E, EMDB 47480).

## Acknowledgements

This work was funded by grants from CIHR (175223) and NSERC (RGPIN-2022-04723) to JEB and by an ERC Starting grant 637733 Pentabrain to HN. CJBD acknowledges support from NSERC (RGPIN-2024-05272). FD acknowledges the State-Region Grand-Est Plan. MJT was supported by an NSERC CGS-D, which included funding to support travel to the IBS in Grenoble, France, for data collection and processing. CJGT was supported by an Ontario Graduate Scholarship. This work used the platforms of the Grenoble Instruct-ERIC center (ISBG; UAR 3518 CNRS-CEA-UGA-EMBL) within the Grenoble Partnership for Structural Biology (PSB), supported by FRISBI (ANR-10-INBS-0005-02) and GRAL, financed within the University Grenoble Alpes graduate school (Ecoles Universitaires de Recherche) CBH-EUR-GS (ANR-17-EURE-0003). The IBS-ISBG EM facility is supported by the Auvergne-Rhône-Alpes Region, the Fondation Recherche Medicale (FRM), the fonds FEDER and the GIS-Infrastructures en Biologie Sante et Agronomie (IBISA). We thank Guy Schoehn for establishing and managing the IBS-ISBG cryo-electron microscopy platform and for providing access, training and support. We also thank Camille Hénault for preparing the nanodisc-reconstituted nAChR samples.

## Author contribution

Initial conceptualization, JEB, HN, EZ, and MJT with subsequent input from CJBD. Data acquisition and processing: ethidium bromide binding and whole cell electrophysiology, MJT and JEB; cryo-EM, MJT, EZ, HN, and JEB: EZ played the primary role acquiring the cryo-EM data sets and supervised the cryo-EM data processing, HN played the primary role supervising model building; MD simulations AA, FD, and JEB; single-channel recording, CJGT, and analysis CJGT, JRE, and CJBD. The initial draft was written by JEB and MJT, with subsequent drafts written by JEB and CJBD, with input from all authors. Figures were prepared by MJT, CJBD, JEB, CJGT, AA, and HN.

## Competing interest declaration

