## Supplemental Figures and Tables for "Asynchronous subunit transitions precede acetylcholine receptor activation"

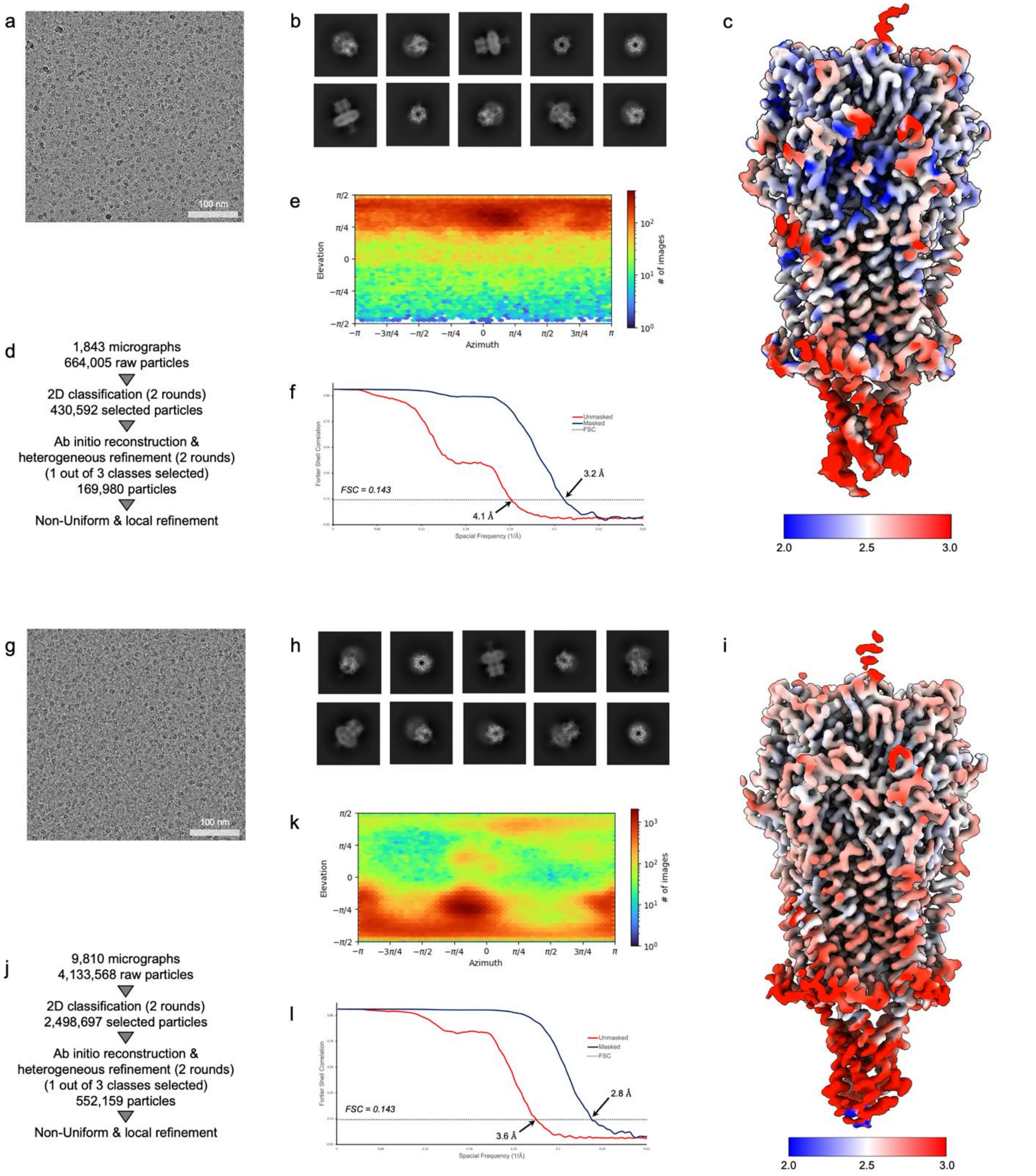


**Figure 1. Data processing for un- and di-liganded states.** Data processing for the un- (panels a-f) and the di- (panels g-l) liganded datasets. Motion and CTF corrected micrographs are shown in panels a and g. Representative 2D projection classifications are shown in panels b and h. Sharpened maps coloured by local resolution are shown in panels c and i. Schematics of the image analysis workflow are shown in panels d and j. Heat maps of the angular distribution of particle projections are shown in panels e and k. Gold-standard Fourier shell correlation (FSC) curves are shown in panels f and l.

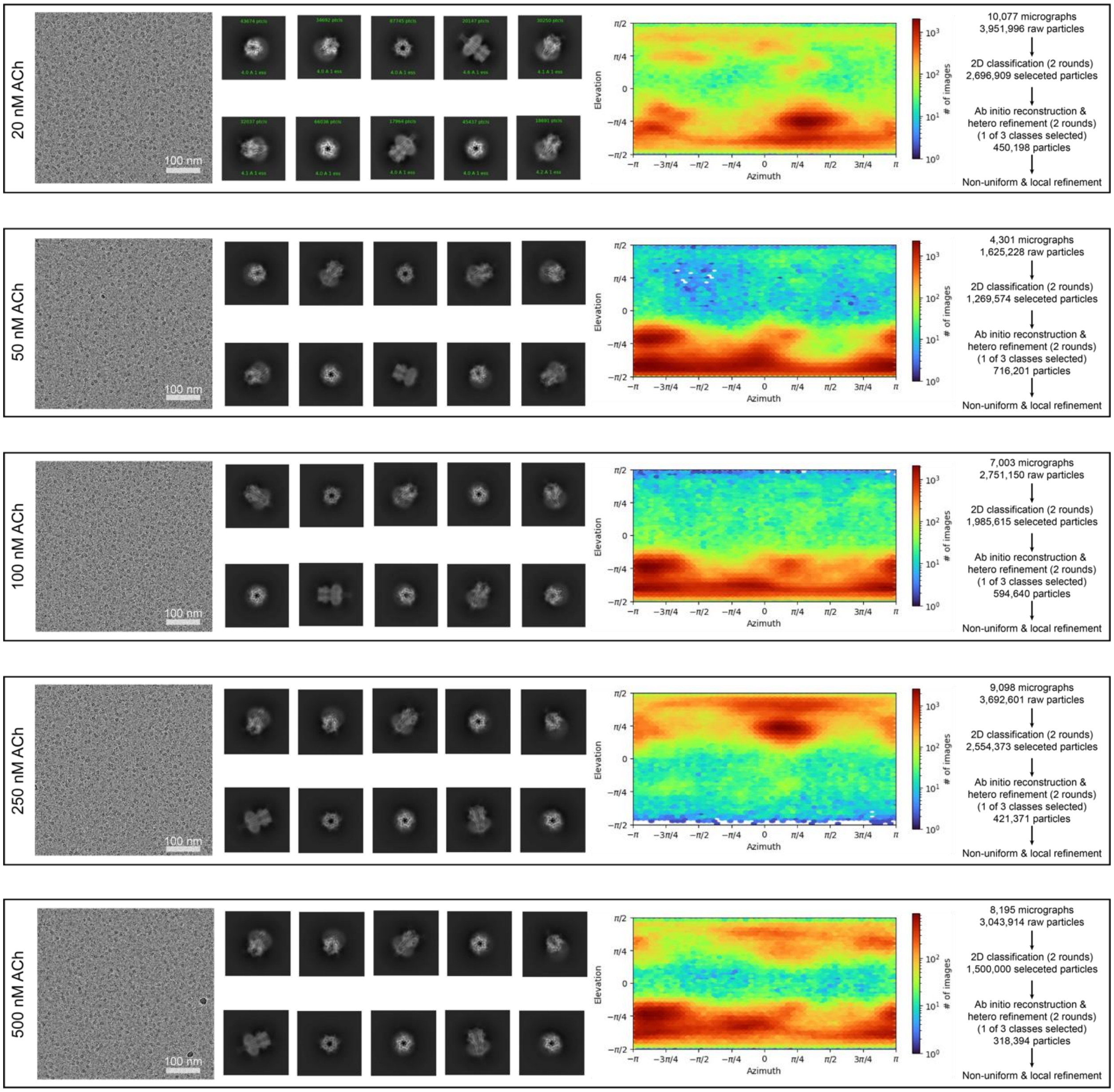


**Figure 2. Data processing for the mono-liganded state.** Datasets at five sub-saturating concentrations of ACh were initially processed individually using the same strategy described for the un- and the di-liganded datasets in Extended Data Fig. 1.

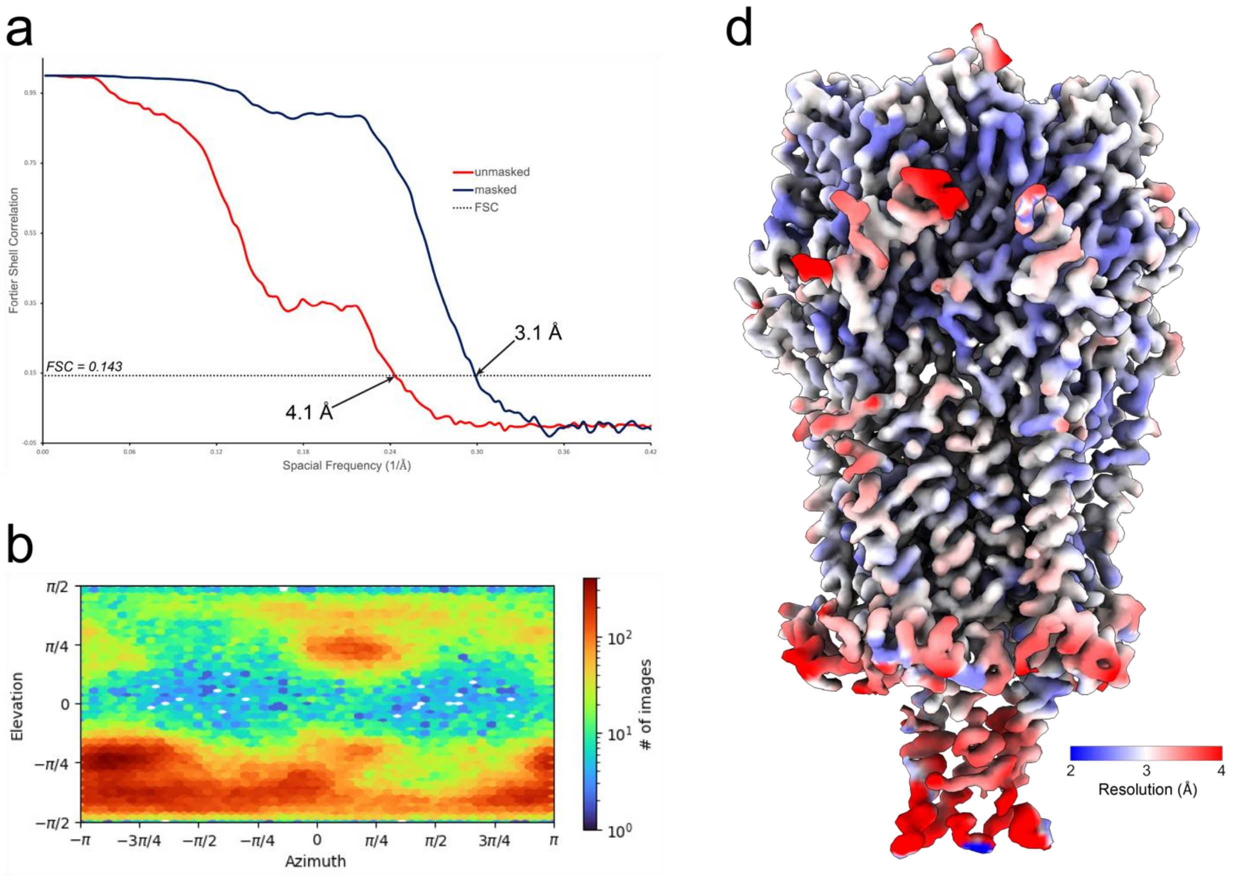


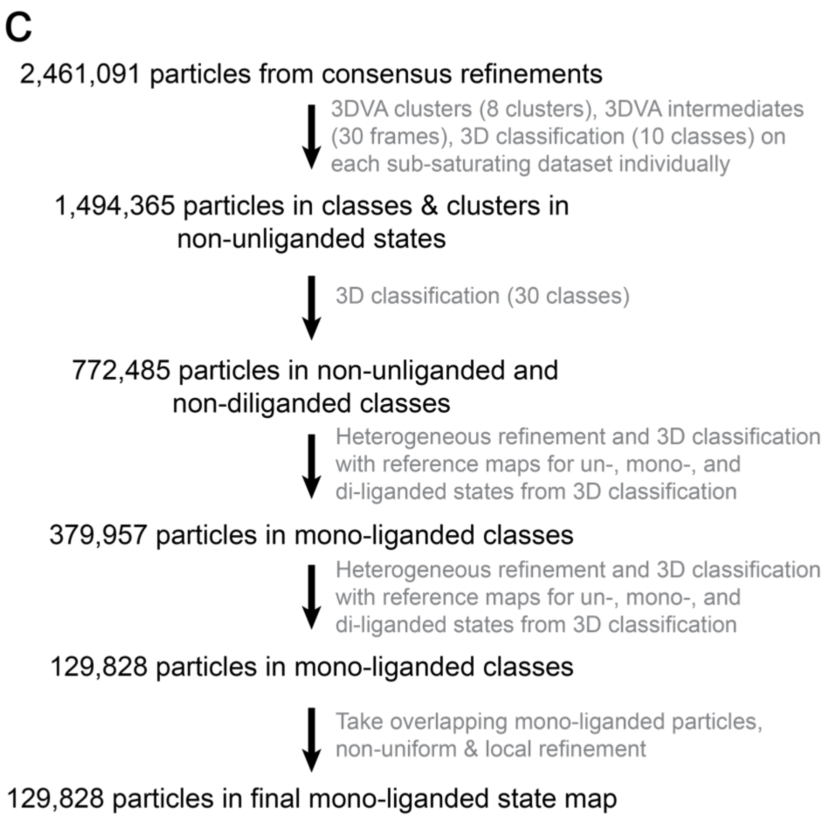


**Figure 3. Quality of the cryo-EM density map for the mono-liganded state.** a) Gold-standard Fourier shell correlation (FSC) curves for the map with and without masking. b) Distribution of particle orientations used in the 3D reconstruction. c) Several rounds of 3D variability analysis and 3D classification were carried out to separate particles corresponding to the mono-liganded state. d) Sharpened map colored by local resolution.

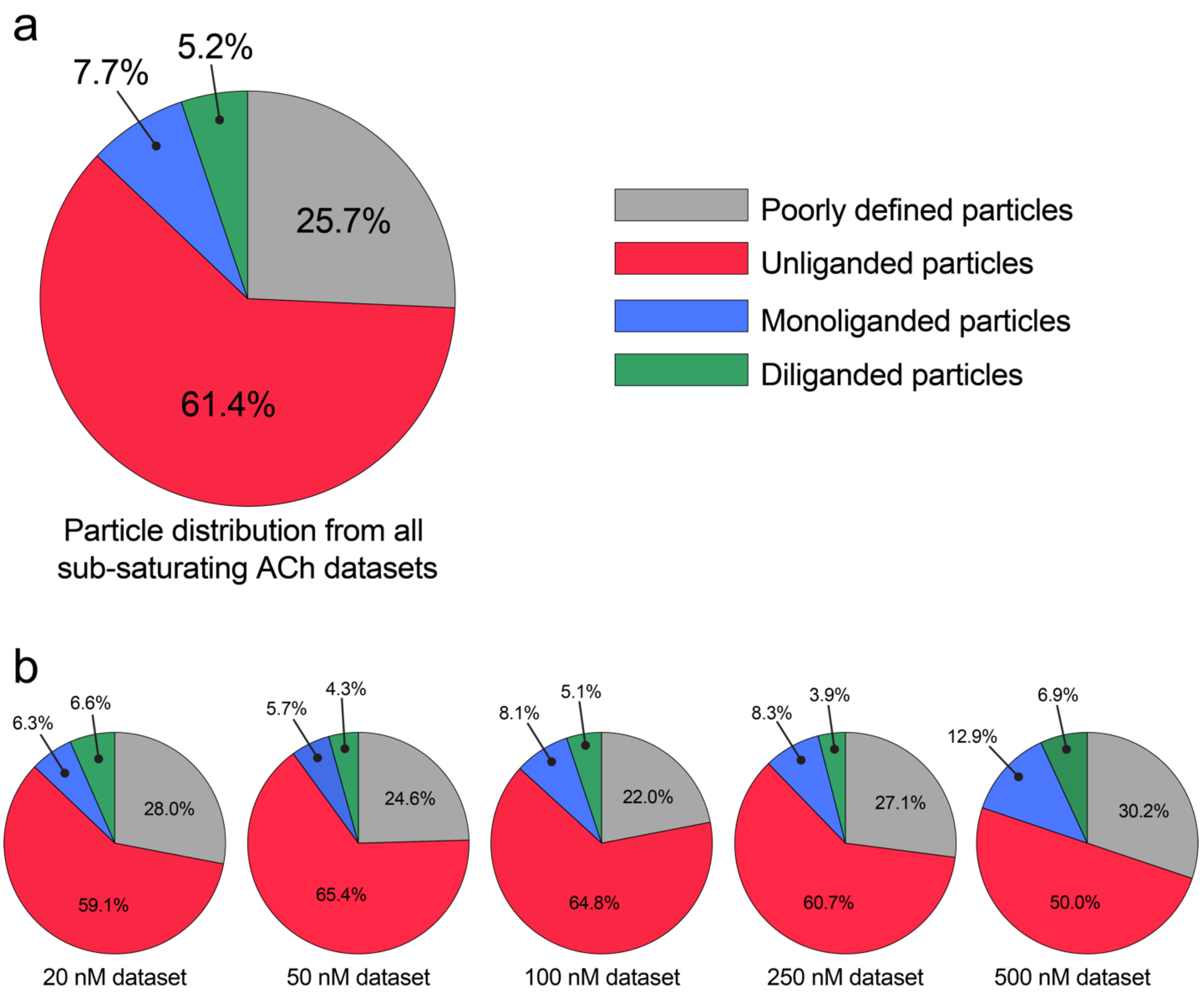


**Figure 4. Particle distributions at sub-saturating ACh concentrations.** (a) Final distribution and (b) individual dataset distributions of particles recorded at sub-saturating concentrations of ACh based on visual interpretation of maps derived from variability and classification analyses. Several maps were difficult to assign due to weak/smeared/fat densities at the binding sites or within the pore. Particles in these maps were discarded and assigned as "poorly defined particles".

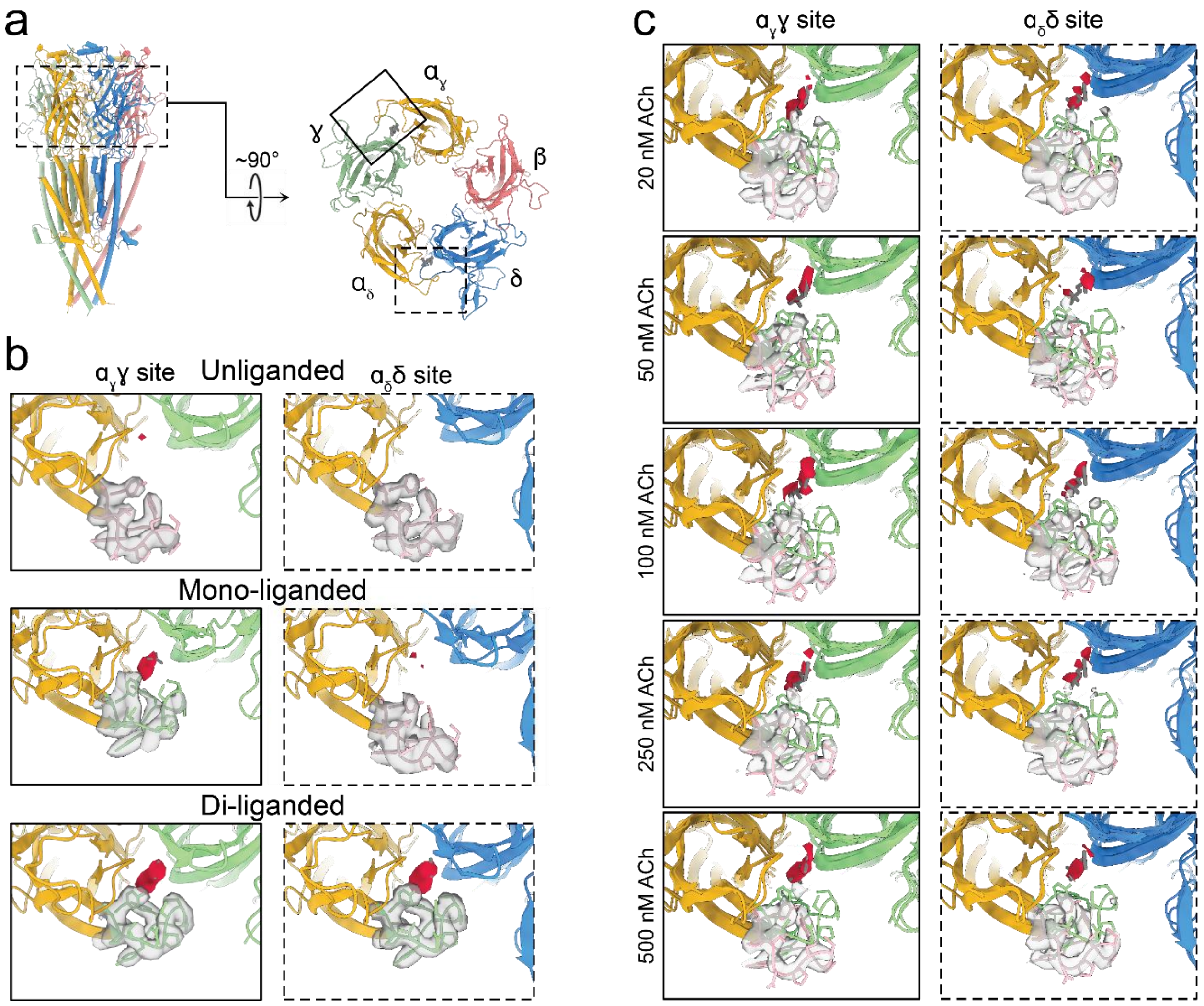


**Figure 5. Binding site density in datasets collected with different concentrations ACh.** a) Side and top views of the nAChR are shown with dashed and undashed boxes depicting the $\alpha_\gamma\gamma$ and $\alpha_\delta\delta$ agonist sites, respectively. b) ACh and loop C density are shown in red opaque and grey transparent surface representations, respectively, with the models for the un-, mono-, and di-liganded states shown in cartoon representation. c) Agonist and loop C density from consensus refinements for all datasets collected at sub-saturating ACh concentrations are shown for both agonist sites. Un- and di-liganded models are shown in cartoon representations with loop C in the unliganded state coloured pink and loop C in the di-liganded state coloured green. Presented densities are from sharpened maps from CryoSparc visualized on ChimeraX. Contour levels were manually adjusted based on density at not variable regions of the nAChR to facilitate comparison. The following contour levels were used: unliganded, 0.1; mono-liganded, 0.2; di-liganded, 0.1; 500 nM, 0.3; 250 nM, 0.35; 100 nM, 0.37; 50 nM, 0.45; 20 nM, 0.3.

**Table 1. Cryo-EM data collection, refinement and validation statistics.**

| | Unliganded | Monoliganded | Diliganded |
|---|---|---|---|
| **Data collection & processing** | | | |
| Microscope | Glacios-IBS | Glacios-IBS | Glacios-IBS |
| Magnification | 36,000 | 36,000 | 36,000 |
| Voltage (kV) | 200 | 200 | 200 |
| Frames (Exposure time in s) | 40 (4) | 40 (4) | 40 (4) |
| Total movies (no.) | 1,843 | 38,674 | 9,810 |
| Electron dose (e/A$^2$) | 42 | 42 | 42 |
| Collection mode | Counting | Counting | Counting |
| Effective pixel size (A) | 1.145 | 1.145 | 1.1 |
| Initial particle images (no.) | 664,005 | 15,064,889* | 4,133,568 |
| Final particle images (no.) | 169,980 | 129,828 | 552,159 |
| Symmetry imposed | C1 | C1 | C1 |
| Map resolution (A) | 3.20 | 3.13 | 2.80 |
| **Model refinement** | | | |
| Initial model used (PDB code) | 7QKO | 7QL5 | 7QL5 |
| **Model composition** | | | |
| Non-hydrogen atoms | 17,181 | 16,980 | 16,853 |
| Protein residues | 2,201 | 2,013 | 2,021 |
| Carbohydrates | 40 | 37 | 36 |
| Lipids | 8 | 5 | 0 |
| **B-factors** (min/max/mean A$^2$) | | | |
| Protein | 101.18/261.86/152.61 | 80.71/416.97/149.15 | 106.36/341.76/154.40 |
| Ligand | 135.24/349.23/194.54 | 110.22/351.13/176.87 | 117.89/316.30/187.46 |
| **R.M.S. Deviations** | | | |
| Bond length (A) | 0.004 (0) | 0.003 (0) | 0.004 (0) |
| Bond angles (0) | 0.566 (3) | 0.521 (2) | 0.540 (1) |
| **Validation** | | | |
| MolProbity score | 1.35 | 1.35 | 1.13 |
| Clashscore | 4.04 | 4.23 | 3.23 |
| Rotamer outliers (%) | 1.02 | 1.24 | 1.02 |
| **Ramachandran plot** (%) | | | |
| Favored | 97.2 | 97.6 | 97.95 |
| Allowed | 2.86 | 2.36 | 2.05 |
| Outliers | 0.00 | 0.00 | 0.00 |
| Model Resolution | 3.8 | 3.7 | 3.4 |

*A total of 2,461,091 particles were used for the consensus refinements in each dataset at sub-saturating ACh and used for down-stream variability analyses.

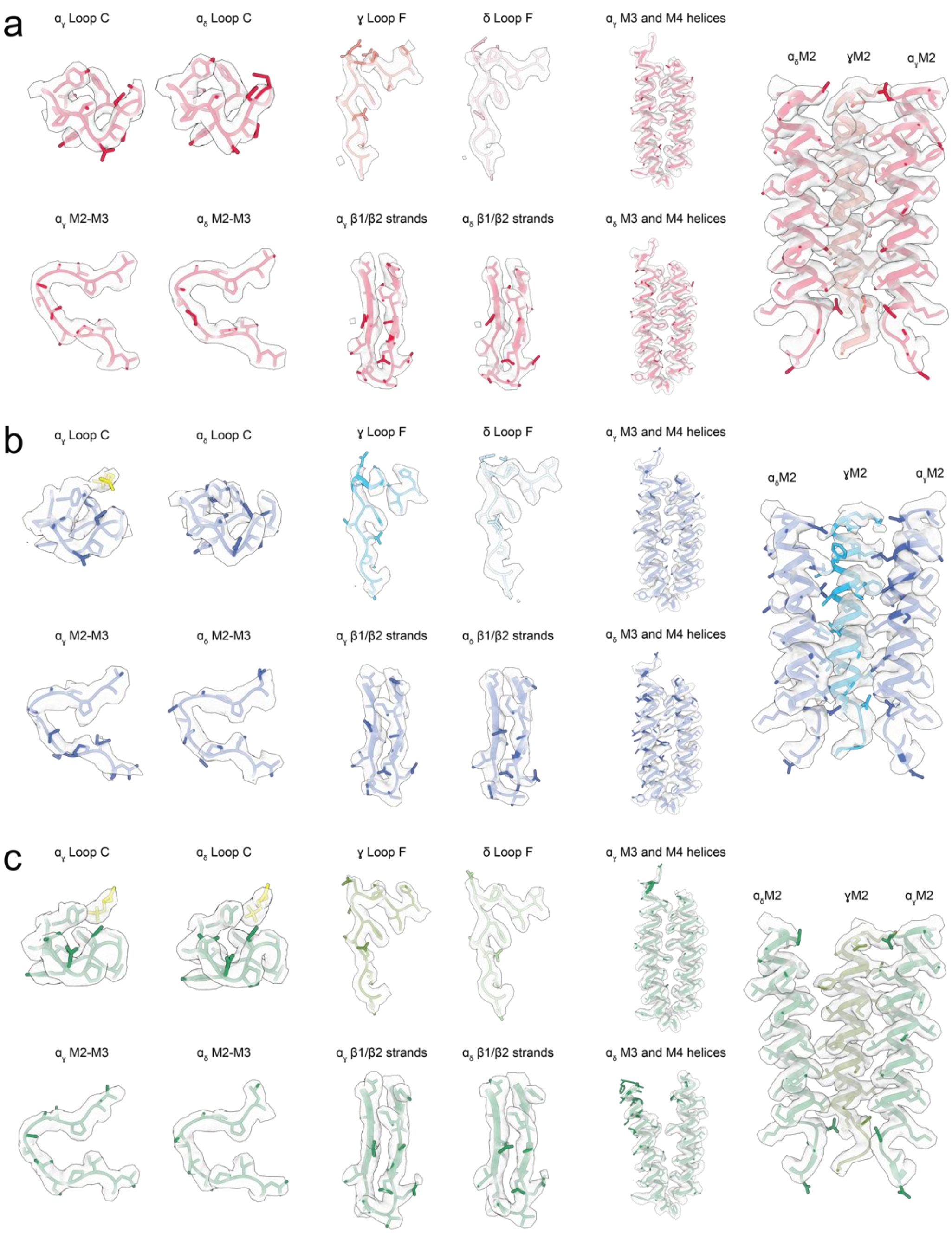


**Figure 6. Quality of the density maps.** Densities in surface representation overlaid on key regions of the unliganded (a), mono-liganded (b), and di- liganded (c) models. Sharpened maps are used for every region except the loop C densities in the mono-liganded state where the unsharpened map was used. The sharpened maps were displayed in ChimeraX at a contour level of 0.1 while the unsharpened map was displayed at a contour level of 0.2.

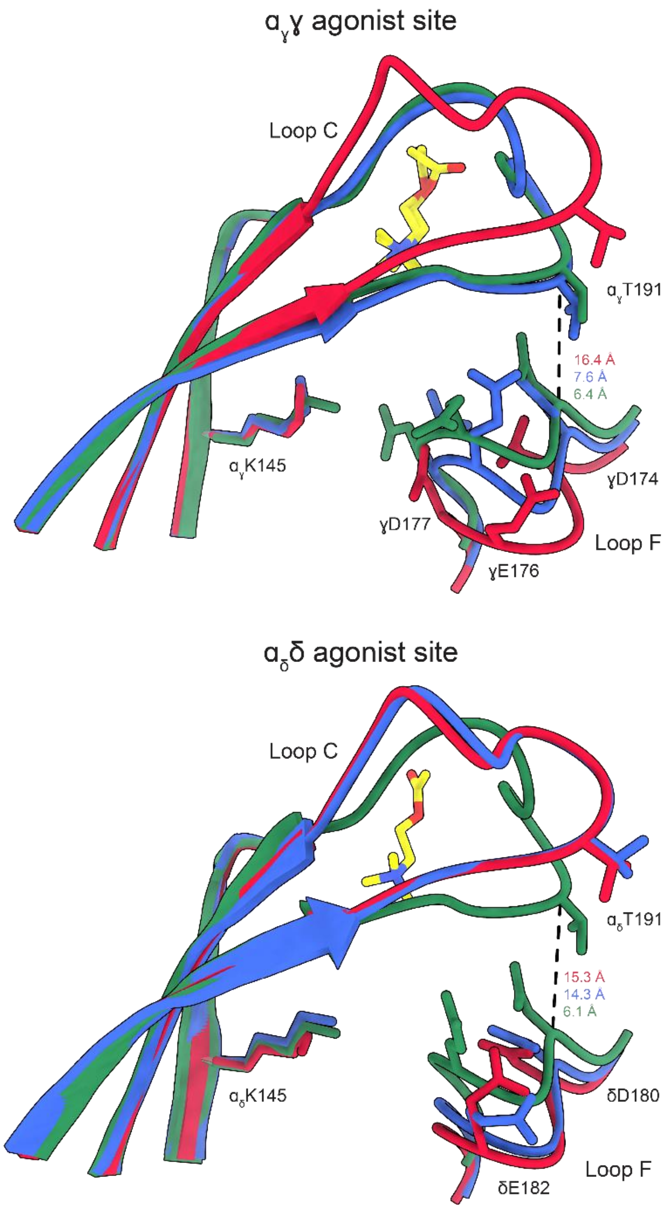


**Figure 7. Loop C and loop F capping in un-, mono- and di-liganded states.** Loop F capping at $\alpha_\gamma\gamma$ (a) and $\alpha_\delta\delta$ (b) sites are shown in unliganded (red), mono-liganded (blue), and di-liganded (green) states. The dashed lines indicate the distance between $C_\alpha$'s from αT191 on loop C and γD174/δD180 on loop F.

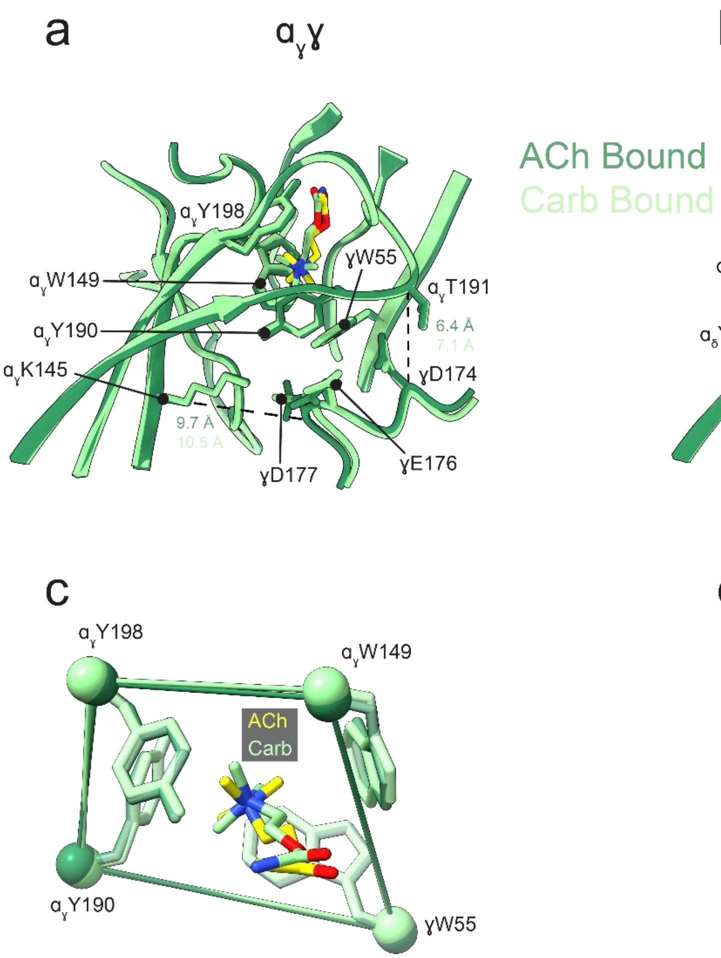


**Figure 8. Comparison of Carb and ACh binding to the nAChR.** Di-liganded ACh bound (dark green) and carbamylcholine (Carb) bound (7QL6, light green) states are superimposed with bound ACh and Carb molecules color yellow and light green, respectively. Interactions between loop F from the complementary γ/δ subunits and the principal α subunits are highlighted in panels a and b for the $\alpha_\gamma\gamma$ and $\alpha_\delta\delta$ agonist binding sites, respectively. The geometry of the aromatic box at the $\alpha_\gamma\gamma$ and $\alpha_\delta\delta$ agonist binding sites are shown in panel c and d, respectively.

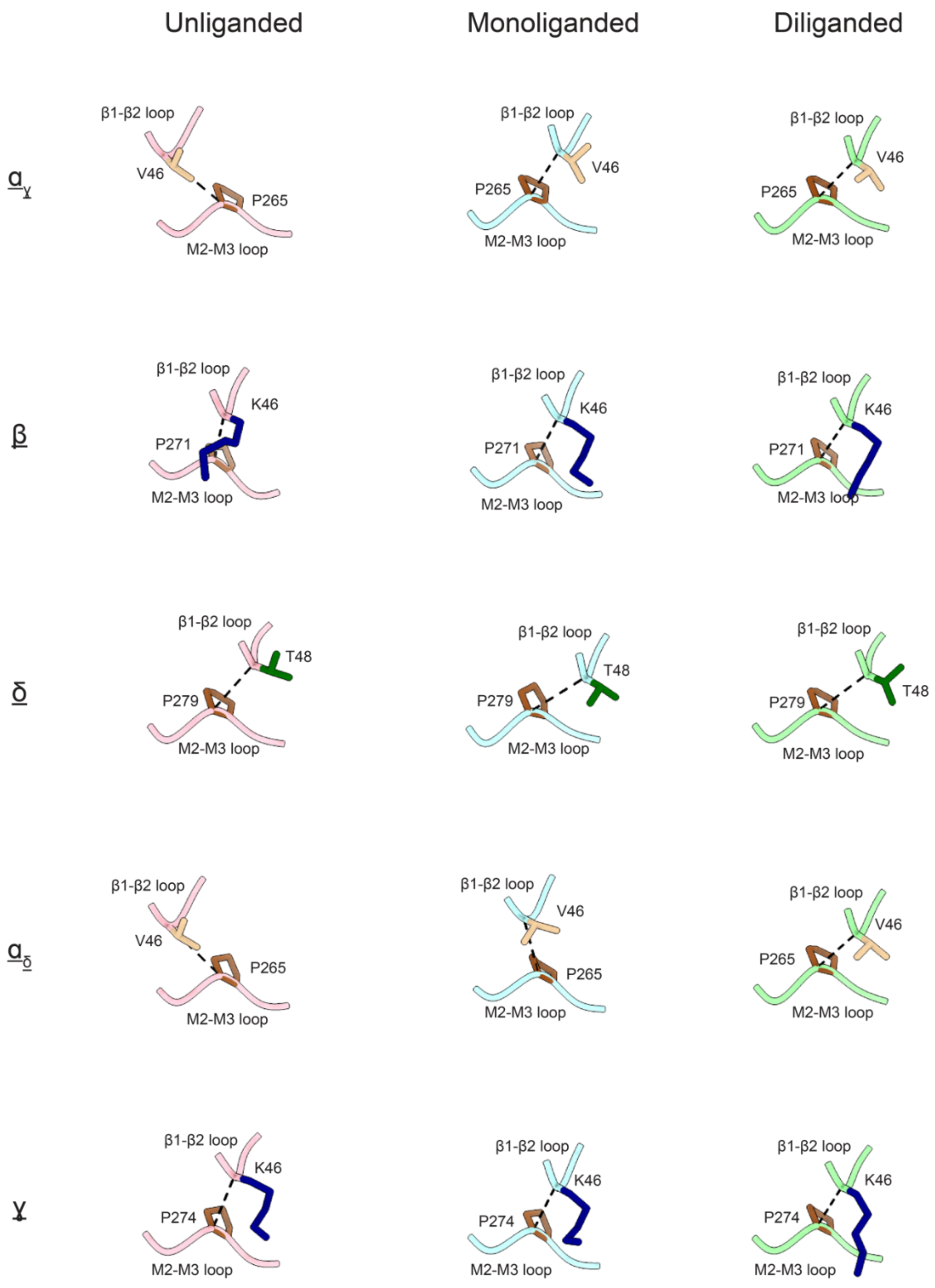


**Figure 9. Relative motions of β1-β2 and M2-M3 loops in each subunit of the un-, mono-, and di-liganded states.** αVal46 and αPro265 (and their equivalent residues in other subunits) are shown to highlight the relative motions that occur at the ECD – TMD interface of each subunit in the three states. While αVal46 transitions from the pore distal side of αPro265 in the unliganded state to the pore proximal side of αPro265 in the di-liganded state, the remaining subunits sit on the pore proximal side throughout. Structures are aligned by their M2-M3 loops to highlight the relative motion of the β1-β2 loop.

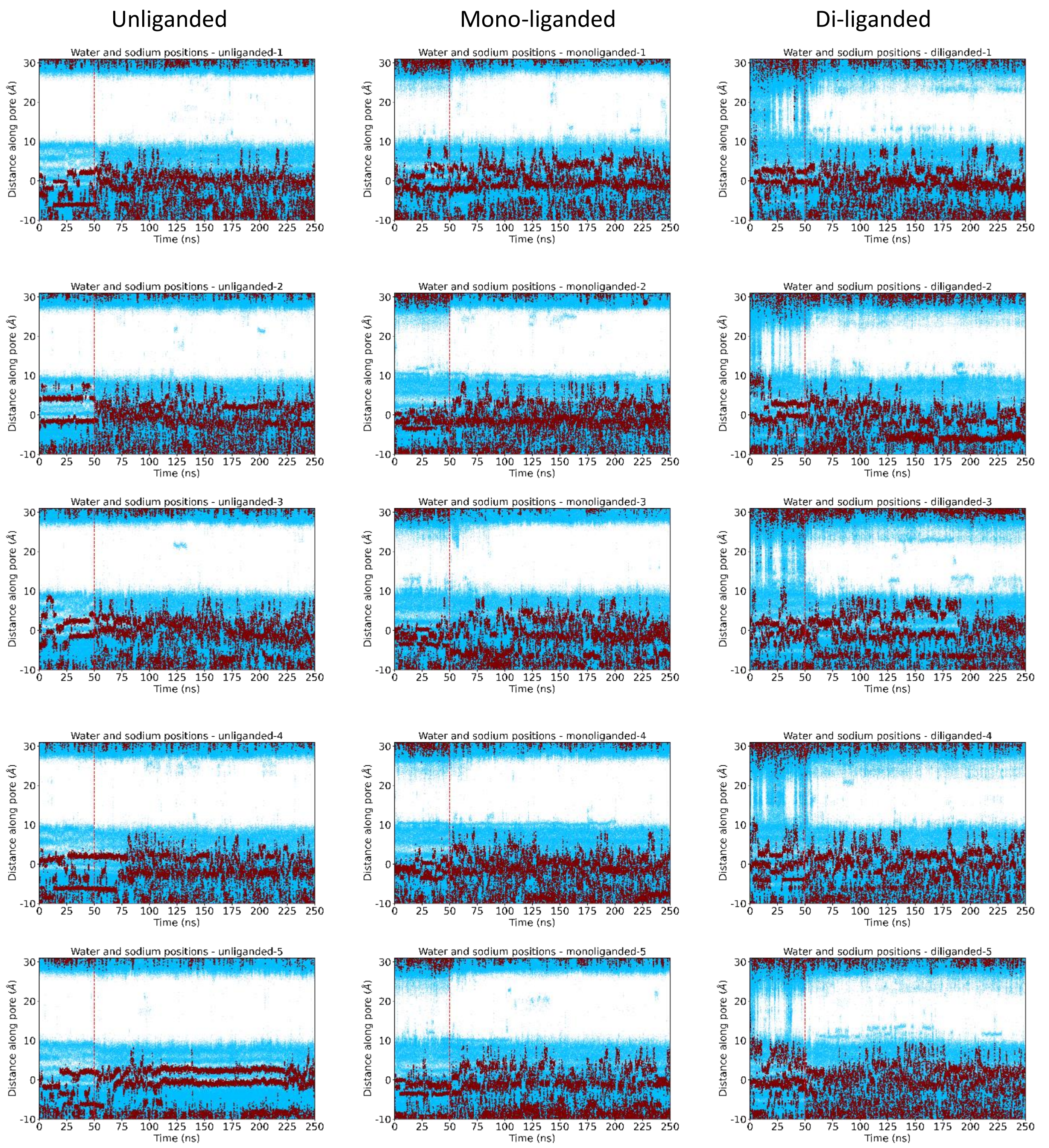


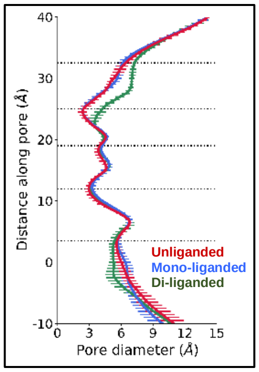


**Figure 10. Hydration of the channel pore in the unliganded, mono-liganded, and di-liganded states. Top five rows:** Trajectories showing water (blue) and sodium (red) ion z coordinates of individual particle positions along the channel axis during molecular dynamics simulation runs, five each for the unliganded, mono-liganded, and di-liganded nAChR structures. Dashed line indicates the release of backbone conformational restraints. **(Left)** Average pore diameters for the three nAChR structures over the final 100 frames of the five trajectories. Error bars represent the standard deviation.

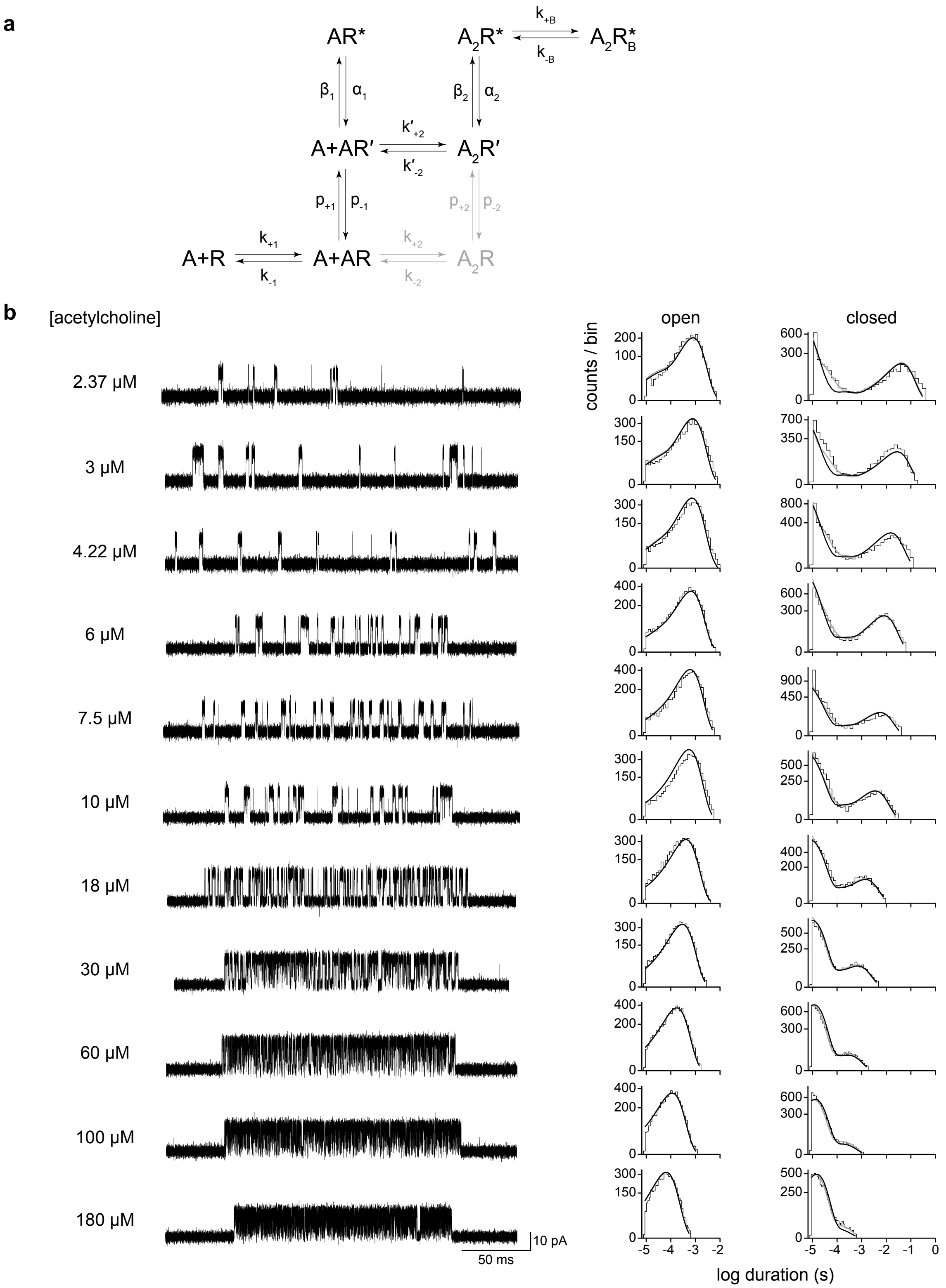


**Figure 11:** Single-channel kinetics of human adult muscle-type acetylcholine receptors. **(a)** Sequential (black) and Prime (includes gray) schemes that include an agonist blocked state flanking the diliganded active state. Estimates of the rate constants for global fits to each scheme can be found in Extended Data, Table 3. Fits were performed on three individual recordings from each presented acetylcholine concentration, from at least two separate transfections, corresponding to a total of 33 patches in the entire global data set. **(b)** Representative bursts (*left*) and corresponding open and closed duration histograms (*right*) from single-channel recordings of the human adult muscle-type acetylcholine receptor over the full acetylcholine concentration range. Global fits at each concentration are ovleraid (sequential scheme = black line; prime scheme = gray line) for the entire data set. Recordings were acquired in the cell-attached patch configuration, and are digitally filtered to 25 kHz, with an applied voltage of –120 mV. Openings are upward deflections, and the scale bar beside the bottom trace represents 50 ms and 10 pA for all traces.

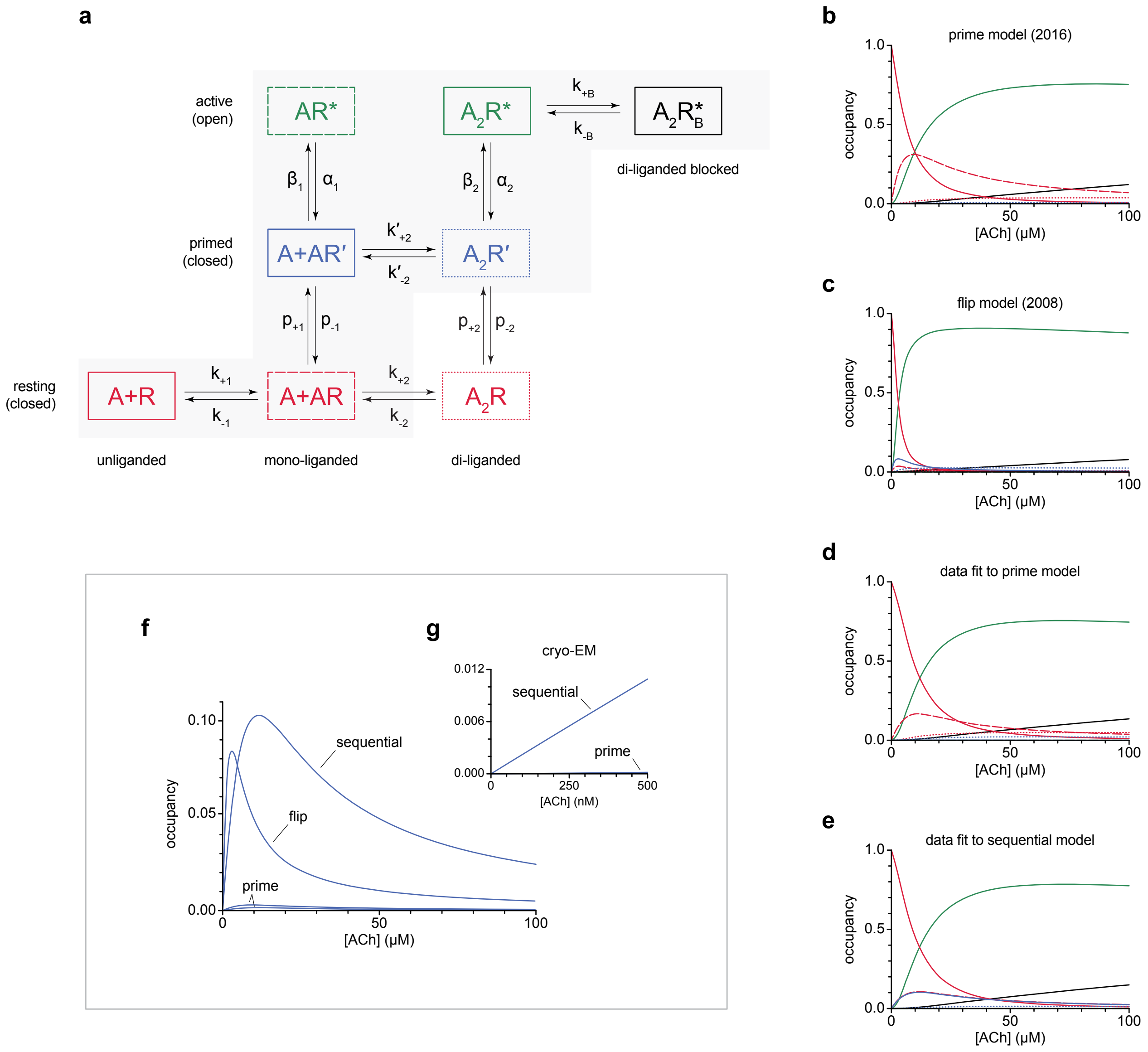


**Figure 12:** A sequential scheme for acetylcholine receptor activation predicts appreciable steady-state occupancy of a mono-liganded intermediate conformation. **(a)** Kinetic schemes of acetylcholine receptor activation that incorporate intermediate ('primed' or 'flipped') states. The sequential scheme used in the present work (gray shaded area) represents a subset of both the prime and flip schemes (all states). Resting states (R) with different levels of agonist (A) occupancy are depicted in red, while primed/intermediate (R´) and active/open (R*) states are blue and green, respectively. Also, included is an agonist blocked state ($R^*_B$) where the channel is open, but current is blocked by agonist (black). **(b-e)** The acetylcholine concentration-dependent steady-state occupancy of every state in each respective scheme can be calculated using rates inferred from global kinetic fitting of single-channel data sets, and were calculated using rates from **(b)** Mukhtasimova *et al.* (2016; prime model ref. 5), **(c)** Lape *et al.* (2008; flip model ref. 3) or the present data fit to **(d)** the prime model or **(e)** the sequential scheme. **(f)** Comparison of the calculated steady state occupancy of the mono-liganded intermediate state predicted by each model shows that the sequential model predicts a peak steady-state occupancy of ~10%, approximating the distribution of particles observed in this state with cryo-EM. **(g)** Calculated steady-state occupancy of the mono-liganded intermediate state over the acetylcholine concentration range used for cryo-EM data collection (includes fits of the presented single-channel data to the prime and sequential models only).

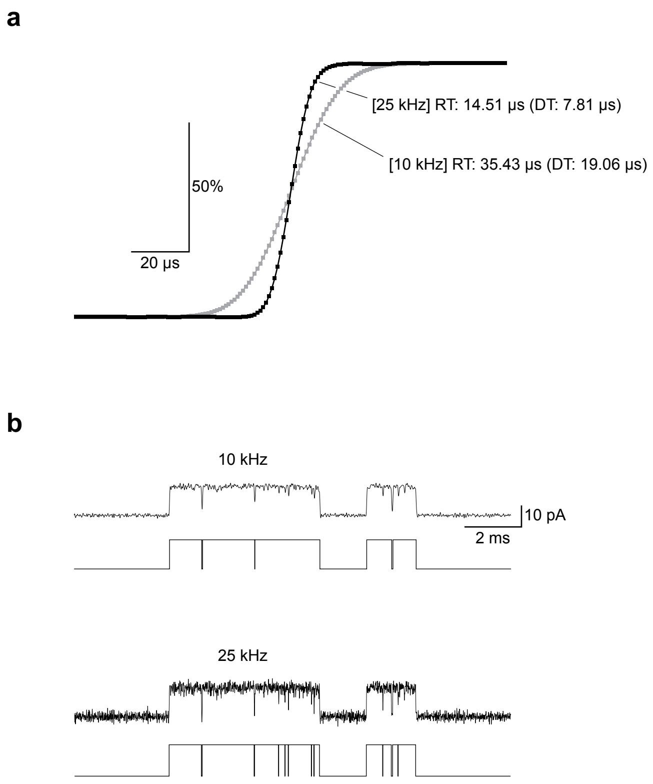


**Figure 13:** High-resolution single-channel recordings make it possible to detect additional brief events. **(a)** Measured step responses for the Axopatch 200B patch clamp amplifier with an internal 100 kHz Bessel filter and a 25 kHz (black), or a 10 kHz (gray) Gaussian digital filter. Data was sampled at 1 μs intervals (data points), and 10,000 individual step responses were aligned and averaged (see Methods). In each case the the 10% to 90% rise time (RT), and resulting dead time (DT; briefest event theoretically detectable) for each condition is indicated. **(b)** Single-channel data obtained from adult human muscle-type AChR at 3 μM acetylcholine with digital Gaussian filter applied at 10 kHz (top) and 25 kHz (bottom). Single-channel openings are presented as upwards deflections. Shown below each filtered trace are the resulting idealized detections obtained using 50% threshold crossings for each filter setting.

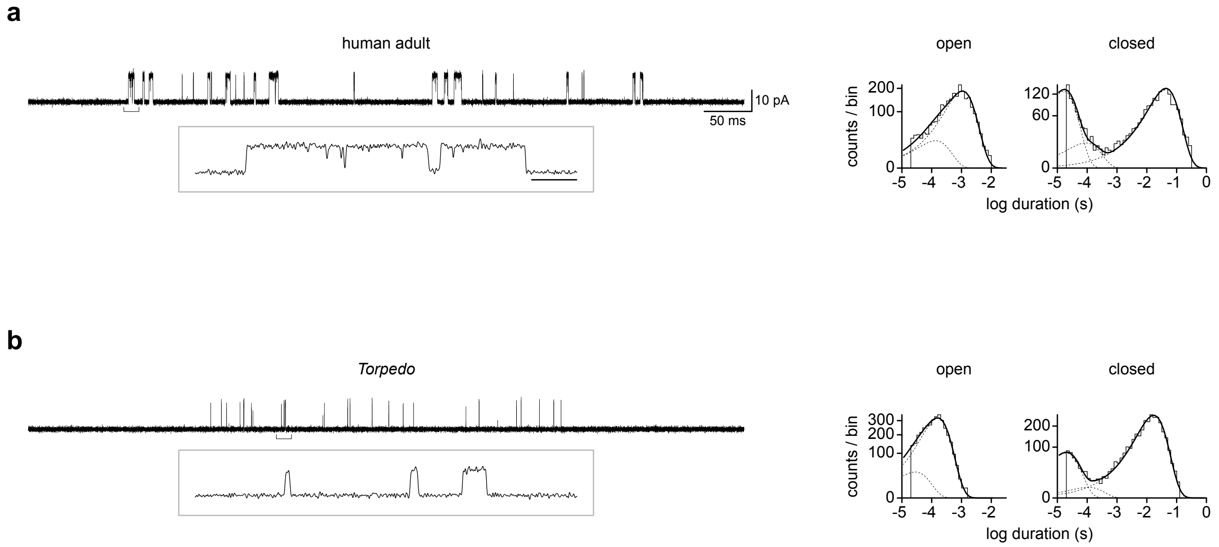


**Figure 14:** The single-channel activity of human and *Torpedo* muscle-type acetylcholine receptors is similar. Bursts of single-channel activity from **(a)** human adult and **(b)** *Torpedo californica* acetylcholine receptors, elicited by 2.37 μM and 10 μM acetylcholine, respectively. Recordings were obtained at –120 mV and digitally filtered to 10 kHz, where openings are upward deflections. Scale bar in 'a' applies to both panels, and the 10 pA vertical bar also applies to the insets. Insets, which depict expansions of the above bracketed areas in each case, were also digitally filtered to 10 kHz. The horizontal scale bar in the inset in 'a' represents 1 ms and also applies to the inset in 'b'. In each case open and closed duration histograms (right) are fit with two or three exponential components (dashed lines), with the sum of exponentials overlaid (solid lines).

**Table 2:** Open duration histogram statistics. Open duration histograms for human adult muscle-type acetylcholine receptors at subsaturating acetylcholine concentrations (20, 100, and 500 nM) were fit with two (left; *i*,*ii*) or three (right; *i*,*ii*,*iii*) exponential components.

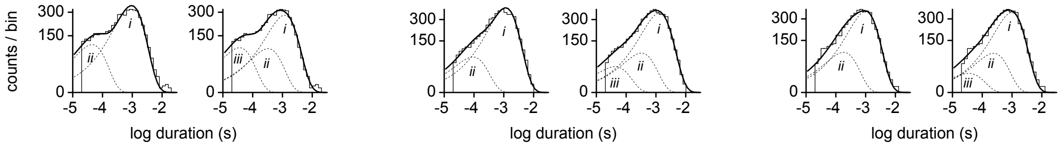


| | **20 nM acetylcholine** | | **100 nM acetylcholine** | | **500 nM acetylcholine** | |
|---|---|---|---|---|---|---|
| | 2 components | 3 components | 2 components | 3 components | 2 components | 3 components |
| log-likelihood(LL) | -47.299 | -23.004 | -20.71 | -10.405 | -23.032 | -17.178 |
| LLR | 24.295 | | 10.305 | | 5.854 | |
| p-value | <0.001 | | <0.001 | | <0.005 | |
| prefered model | No | Yes | No | Yes | Yes | No |
| SIC score | 64.29 | 48.49 | 37.89 | 36.18 | 40.13 | 42.83 |
| ranking | 2 | 1 | 2 | 1 | 1 | 2 |
| prefered model | No | Yes | No | Yes | Yes | No |

***Note:*** Log-likelihoods (LL) were calculated within TACFit for each fit of the histograms (see Methods). Log-likelihood ratio (LLR) tests and Schwarz information criterion (SIC) were used to evaluate the need for the 3rd component in each of the histograms. LLRs were calculated according to:

$$LLR = LL_{k1} - LL_{k2}$$

Where 'k1' is the model with more free parameters (in this case three exponential components), and 'k2' the model with fewer free parameters (in this case two exponential components). SIC scores were calculated as:

$$SIC = -LL + 0.5F*\ln(N)$$

Where 'F' is the number of free parameters and 'N' the number of open dwells included in the histogram. Models are ranked higher if their SIC is lower.

**Table 3:** Inferred rate constants from maximum likelihood kinetic fitting. Rate constants were estimated by fitting sequential (black) and primed (includes gray) kinetic schemes to a global data set spanning a full range of acetylcholine concentrations. Associated errors were computed within MIL, and are presented in parentheses.

| | $^*k_{+1}$ | $k_{-1}$ | $K_1$ | $k_{+2}$ | $k_{-2}$ | $K_2$ | $^*k'_{+2}$ | $k'_{-2}$ | $K'_2$ | $p_{+1}$ | $p_{-1}$ | $P_1$ |
|---|---|---|---|---|---|---|---|---|---|---|---|---|
| sequential | $3.30\times10^{8*}$ | $1.43\times10^{4}$ | 43.3 | | | | $3.20\times10^{8*}$ | $5.50\times10^{4}$ | 17.2 | $2.61\times10^{4}$ | $2.70\times10^{4}$ | 0.967 |
| | $(2.0\times10^{1})$ | $(1.80\times10^{3})$ | | | | | $(3.0\times10^{1})$ | $(5.70\times10^{2})$ | | $(1.45\times10^{3})$ | $(4.00\times10^{3})$ | |
| prime | $3.60\times10^{8*}$ | $1.02\times10^{4}$ | 28.3 | $4.10\times10^{8*}$ | $3.20\times10^{4}$ | $7.8\times10^{7}$ | $1.0\times10^{9*}$ | $1.70\times10^{3}$ | 1.7 | $2.50\times10^{2}$ | $2.63\times10^{4}$ | 0.010 |
| | $(1.50\times10^{1})$ | $(9.00\times10^{2})$ | | $(2.0\times10^{1})$ | $(1.10\times10^{3})$ | | $(1.30\times10^{2})$ | $(1.70\times10^{2})$ | | $(2.0\times10^{1})$ | $(2.25\times10^{3})$ | |

| | $p_{+2}$ | $p_{-2}$ | $P_2$ | $\beta_1$ | $\alpha_1$ | $\Theta_1$ | $\beta_2$ | $\alpha_2$ | $\Theta_2$ | $k_{+B}$ | $k_{-B}$ | $K_B$ |
|---|---|---|---|---|---|---|---|---|---|---|---|---|
| sequential | | | | $1.00\times10^{2}$ | $4.50\times10^{4}$ | 0.0022 | $^{a}1.64\times10^{5}$ | $3.00\times10^{3}$ | 54.7 | $1.75\times10^{2*}$ | $9.02\times10^{4}$ | 0.002 |
| | | | | $(1.0\times10^{1})$ | $(3.4\times10^{3})$ | | $(1.25\times10^{4})$ | $(2.00\times10^{2})$ | | $(1.00\times10^{0})$ | $(8.50\times10^{2})$ | |
| prime | $1.65\times10^{4}$ | $3.80\times10^{4}$ | 0.434 | $8.80\times10^{3}$ | $4.65\times10^{4}$ | 0.1892 | $^{a}4.70\times10^{5}$ | $1.33\times10^{4}$ | 35.3 | $2.0\times10^{2*}$ | $1.1\times10^{5}$ | 0.002 |
| | $(7.0\times10^{2})$ | $(2.0\times10^{3})$ | | $(1.1\times10^{3})$ | $(3.3\times10^{3})$ | | $(1.20\times10^{4})$ | $(9.00\times10^{2})$ | | $(1.50\times10^{0})$ | $(7.50\times10^{2})$ | |

***Note:*** Units are $M^{-1}s^{-1}$ for association rate constants(*), and $s^{-1}$ for all other rate constants. Equilibrium dissociation constants are presented in μM. Equilibrium constants were calculated as follows:

$$K_n = k_{-n}/k_{+n}$$
$$P_n = p_{+n}/p_{-n}$$
$$\Theta_n = \beta_n/\alpha_n$$

****Note:*** Rates that are dependent upon the acetylcholine concentration.

***ᵃNote:*** While $\beta_2$ appears large, it is likely inflated for the same reasons discussed previously [refs. 5,30].

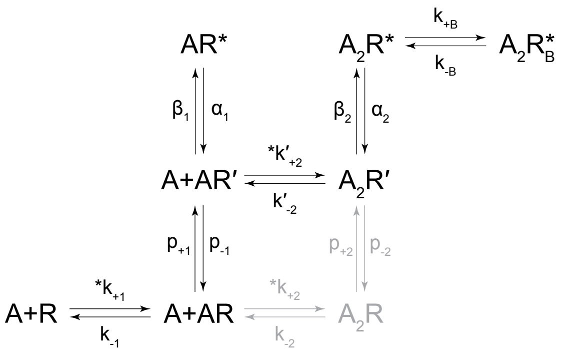


schemes

**Table 4:** Recording statistics for single-channel data included in kinetic fitting. Recordings were performed in triplicate, with the total number of events for each acetylcholine concentration being the sum of the events for each of the three individual recordings. Bursts were classified based on a $\tau_{crit}$, which was specific to each recording (see Methods). Mean $P_{open}$ and standard deviations (SD) were calculated from all bursts included in the analysis (see Methods).

| acetylcholine concentration (μM) | no. of recordings | no. of events | no. of bursts | mean $P_{open}$ | $P_{open}$ (SD) |
|---|---|---|---|---|---|
| 2.37 | 3 | 16635 | 196 | 0.031 | 0.014 |
| 3 | 3 | 25787 | 285 | 0.083 | 0.028 |
| 4.22 | 3 | 24621 | 151 | 0.097 | 0.042 |
| 6 | 3 | 30419 | 141 | 0.182 | 0.048 |
| 7.5 | 3 | 28053 | 233 | 0.266 | 0.087 |
| 10 | 3 | 29279 | 116 | 0.316 | 0.083 |
| 18 | 3 | 32033 | 59 | 0.567 | 0.065 |
| 30 | 3 | 31935 | 60 | 0.700 | 0.072 |
| 60 | 3 | 31455 | 25 | 0.757 | 0.028 |
| 100 | 3 | 31959 | 34 | 0.769 | 0.037 |
| 180 | 3 | 33121 | 37 | 0.726 | 0.027 |

**Table 5:** Model statistics. Depicted kinetic schemes were fit to the same sequence of single-channel dwells in the global data set, which spanned eleven different concentrations of acetylcholine (from 2.37 to 180 μM; see Extended Data, Table 4) that all elicited clear bursts of single-channel activity.

| | **sequential model** | **prime model** |
|---|---|---|
| log-likelihood (LL) | 2409014.5 | 2411895.75 |
| number of events | 313993 | 313993 |
| number of free parameters | 12 | 15 |
| LLR | 2881.25 | |
| p-value | 0 | |
| preferred model | no | yes |
| SIC | -2408939.6 | -2411800.8 |
| ranking | 2 | 1 |
| preferred model | no | yes |
| kinetic scheme | 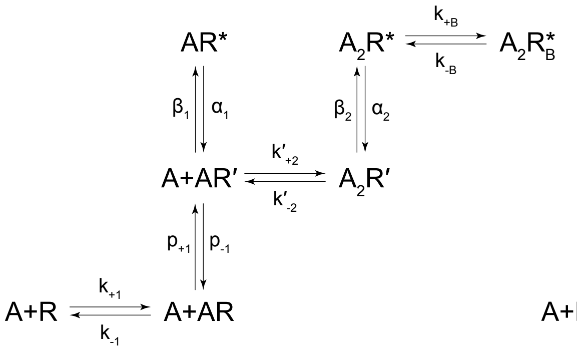 | 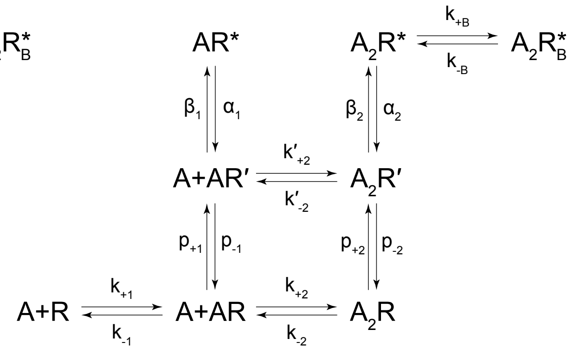 |





***Note:*** Models were compared using their log likelihoods (LL) obtained within the program MIL (see Methods). Log likelihood ratio tests (LLR) and the Schwarz information criterion (SIC) were used to compare how well the two models fit the experimental data. LLRs were calculated according to:

$$LLR = LL_{k1} - LL_{k2}$$

Where ‘k1’ is the model with more free parameters (prime), and ‘k2’ the model with fewer free parameters (sequential). SIC scores were calculated as:

$$SIC = -LL + 0.5F*\ln(N)$$

Where ‘F’ is the number of free parameters and ‘N’ is the number of dwells included in the global data set. The model with the smallest SIC score is the highest ranking.
